# Genetic basis of immunity in Indian cattle as revealed by comparative analysis of *Bos* genome

**DOI:** 10.1101/2024.12.09.627532

**Authors:** Menaka Thambiraja, Shukrruthi K. Iyengar, Brintha Satishkumar, Sai Rohith Kavuru, Aakanksha Katari, Dheer Singh, Suneel K. Onteru, Ragothaman M. Yennamalli

**Affiliations:** Department of Bioinformatics, School of Chemical and Biotechnology, SASTRA Deemed to be University, Thanjavur, India; Molecular Endocrinology, Functional Genomics and Systems Biology Laboratory, Animal Biochemistry Division, ICAR-National Dairy Research Institute, Karnal, India

**Author notes:** Co-Corresponding Authors, Dr. Suneel K. Onteru, Molecular Endocrinology, Functional Genomics and Systems Biology Laboratory, Animal Biochemistry Division, ICAR-National Dairy Research Institute, Karnal, India 132001., Ph no, Dr. Ragothaman M. Yennamalli, Department of Bioinformatics, School of Chemical and Biotechnology, SASTRA Deemed to be University, Thanjavur, Tamil Nadu 613401, India., Ph no. Equal contribution (Shukrruthi K. Iyengar, Brintha Satishkumar, Sai Rohith Kavuru, Aakanksha Katari).

## Abstract

Indicine cattle (*Bos indicus*) show notable resilience and disease resistance compared with taurine breeds, but the genomic basis of these traits remains largely unexplored. Identification of genomic elements for immunity will enable future controlled crossbreeding programs using molecular breeding methods. Therefore, we performed a genome-wide comparison among Nelore, Gir, and Hereford breeds using their whole-genome sequences, majorly focusing on immune-related structural and sequence variation. Our aims were to catalog insertions, deletions, and single nucleotide variants (SNVs) that intersect immune loci and known quantitative trait loci (QTLs), identify runs of homozygosity and selective-sweep signals, and prioritize candidate genes for follow-up functional studies. We retrieved whole-genome sequencing data for Nelore breed (n=14) and Gir breed (n=20) from NCBI using the SRA toolkit. Reads were checked with FastQC, filtered with fastp to remove low-quality bases and adaptors, and retained high-quality reads based on Q20 and Q30. The reads were mapped to the *Bos taurus* reference (ARS-UCD2.0) with BWA-MEM; alignments were processed with SAMtools for sorting, duplicate marking, and MAPQ ≥ 20 filtering. Variants (insertions, deletions, SNVs) were called with GATK HaplotypeCaller, hard-filtered, normalized with bcftools, and annotated with SnpEff and SnpSift. Common variants were identified via in-house Python scripts; immune loci were detected from InnateDB and keyword searches; QTL overlaps were identified using Animal QTLdb; DAVID was used for GO and KEGG enrichment (P < 0.05). ROH islands were defined in PLINK as regions shared by >50% of individuals or samples, and selective sweeps were detected with RAiSD; genes overlapping ROH islands and RAiSD peaks were prioritized as candidate selection signatures. GATK identified 1,884,058 indels and 13,997,533 SNVs in Nelore breed, and 1,457,337 indels and 11,627,881 SNVs in Gir breed, with Ti/Tv ratios of ∼2.26 and ∼2.25, respectively. Nelore breed has more number of variants than Gir. We observed frameshift insertions in *TLR3* and *LOC508441 (CD33)* in both the breeds and frameshift deletions in *JAM3* in Nelore breed and *PAX5* in Gir breed. The variants are also identified in the regulatory regions of both breeds. The high-impact SNVs were in *CD46* and *IL26* genes in Nelore breed, and *PLG* gene in Gir breed. Genome-wide scans using RAiSD identified selective sweeps in 707 candidate genes in Nelore breed and 165 in Gir breed. Comparing the ROH and RAiSD results, we prioritized the genes *ANKRD11, MAGI2, LOC132345096, FOXP2, TCF12*, and *ATP5PO* in Nelore breed, and the genes *MEFV* and *ORIF1* in Gir breed. These genes are found in QTLs linked to milk and health traits. Functional enrichment showed that the genes exhibiting all the three variants belong to immune pathways such as, NF-kappaB signaling, T-cell receptor signaling, and MAPK signaling in both breeds. These results reveal breed-specific genomic variation locating immune loci and its associated QTLs and provide a list of candidate genes and regions for experimental validation and marker development to improve disease resistance and productivity in Indicine cattle. **Keywords:** Hereford, Nelore, Gir, immune-related genes, QTLs, Whole genome sequence

## Introduction

Indicine cattle (members of *Bos indicus*), such as zebu cattle, are native to the Indian subcontinent. Accounting for about 80% of India’s total cattle population, they offer various economic advantages over taurine cattle (members of *Bos taurus*), particularly when raised under tropical conditions [1, 2]. The native breeds are crucial to the country’s dairy industry, which has made India a global leader in milk production, and supports over 80 million rural households. Currently, 50 distinct breeds of indicine cattle are registered in India, evolved through both natural selection and selective breeding for adaptation to different climatic conditions. These breeds can cope with extreme heat or similar adverse climatic conditions, survive despite poor nutrition, and are resistant to several diseases [3, 4, 5, 6]. One notable example is ‘Vechur’ (named after a village of the same name), the smallest cattle breed in the world, found in the Kottayam district of Kerala. It is well-known for its resistance to viral, bacterial, and parasitic diseases than that shown by other exotic breeds and crossbred cattle [7].

The innate immune response, which kicks in at the earliest stages of infection in mammals, protects them from pathogenic infections. This response is non-specific to the pathogen [8, 9, 10]. Therefore, understanding the differences between the immune systems of indicine and taurine cattle could provide valuable insights into disease resistance shown by members of *Bos indicus.* This disease resistance trait is essential for better management of herds and breeding development strategies [11, 12]. In particular, exploring the genomic regions responsible for disease resistance and susceptibility variations opens new opportunities to identify significant genomic signatures for future molecular breeding studies [13].

Recent research on livestock improvement is increasingly focused on whole-genome data to detect genetic factors or genomic signatures associated with important traits such as disease resistance. This includes identifying such structural variations such as insertions, deletions, duplications, inversions, translocations, and other structural variations present on genomic regions by comparing indicine and taurine breeds. Two relevant sources of assembled whole-genome data at chromosome level of cattle are (1) the first whole-genome sequence of *Bos indicus* cattle (Nelore breed) with 52X coverage by the SOLiD sequencing platform with a short-read length of 25–50 bases [14] and (2) the whole-genome sequencing data of *Bos taurus* cattle with multiple improvements, such as covering 91 % of the genome, closing gaps, correcting errors, removing bacterial contigs, and identifying the first portion of Y chromosomes using whole-genome shotgun sequencing methods [15]. Recently, a comparative whole-genome analysis of several *Bos indicus* breeds (Kangayam, Tharparkar, Sahiwal, Red Sindhi, and Hariana) and of *Bos taurus* cattle identified a number of genetic variations. This study revealed over 155 million single-nucleotide polymorphisms (SNPs) and more than one million InDels among a total of 17,252 genes [16]. Among these genes, many involved in innate immune responses were associated with key pathways such as toll-like receptor signaling, retinoic acid-inducible gene I-like receptor signaling, NOD-like receptor signaling, and the Jak-STAT pathway. The same study identified many missense variants in genes such as *TLR3* and *TLR4*, transcription factors such as IRFs (IRF3/IRF7) and NF-kB, and non-synonymous variants in genes such as *MyD88, IRAK4, RIG-I, TRIM25, MAVS, NOD1*, and *NOD2* [16]. Furthermore, draft *de novo* genome assemblies and mitochondrial genome assemblies of *Bos indicus* breeds such as Ongole, Kasargod Dwarf, Kasargod Kapila, and Vechur were generated using Illumina short-read technology. An analysis of the exon-intron structure of 15 key genes related to bovine traits including milk quality, metabolism, and immune response revealed structural variation in their exon-intron numbers. Notably, seven of these genes had fewer exons in Ongole than in *Bos taurus*. Among those genes, *PFKP* and *GPX4* are involved in glycolysis regulation and immune response [17].

Over the past decade, several studies have explored the genetic basis of economically important traits, including disease resistance, in cattle using genome-wide data. For instance, Koufariotis et al. examined whole-genome sequence variants in coding and regulatory regions to evaluate their contribution to dairy traits, highlighting the role of functional annotations in understanding trait architecture [18]. Similarly, Álvarez Cecco et al. performed a genome-wide selection scan in Brangus cattle, uncovering genomic regions likely shaped by artificial and natural selection in composite breeds [19]. Ancestral allele analysis, as demonstrated by Naji et al. and expanded further by Dorji et al., has provided evolutionary insights into bovine genomic variation by defining ancestral states across millions of variants [20,21]. Utsunomiya et al. integrated different genome-wide selection scan methods to detect loci under recent positive selection in dairy and beef cattle, identifying potential targets of selection related to production and adaptation traits [22].

Although *Bos indicus* assemblies are available in NCBI, their utility for applied genetic analyses is highly limited due to quality concerns. The Nelore assembly (*Bos_indicus*_1.0, RefSeq: GCF_000247795.1) has been flagged by NCBI as RefSeq annotation is surpassed [14]. The Gir assembly (ASM293397v1) is reported as contaminant as well unannotated. The assembly methods for the Gir genome are unpublished as of yet. These shortcomings restrict their reliability for comparative or functional studies, especially when contrasted with the high-quality *Bos taurus* reference genome (Hereford, ARS-UCD2.0, RefSeq: GCF_002263795.3), which is widely accepted by the scientific community. The limitations can be overcome by analyzing other high-quality whole-genome sequences together, which is the focus of this study.

Despite the multiple studies exploring the genetic basis of various traits in cattle, a genome-wide comparative study in the focus of immunity in *Bos indicus* breed remains largely unexplored. The crossbreeding program in India focused on enhancing the milk production. However, the crossbreds are observed to be susceptible to various diseases compared to other Indicine cattle. Hence, there is a need to identify the genomic elements in the Indicine genome that can contribute towards innate immunity. Thereby, future controlled crossbreeding programs can be performed using molecular breeding methods to induce immunity against prevalent diseases. Apart from innate immunity genes, the present study also focused the gene’s association with QTLs related to traits such as milk production, reproduction, health, meat and carcass, and exterior QTLs. Among the many Indicine cattle, Nelore and Gir breed sequence data were available for performing whole genome sequence comparison studies. Therefore, we performed detailed study on the genomic variations located in the immune-related genes in Nelore and Gir breed.

## Materials and methods

### General workflow

We retrieved whole-genome sequencing data for Nelore breed (n = 14) and Gir breed (n = 20) from NCBI using the SRA Toolkit (https://trace.ncbi.nlm.nih.gov/Traces/sra/sra.cgi?view=software). The raw read’s quality was checked with FastQC [23], filtering with fastp [24] to remove low-quality bases and adaptor contaminants, and high quality reads based on Q20 and Q30 scores were retained. BWA-MEM [25] was used to map the reads to the *Bos taurus* reference genome (ARS-UCD2.0) and processed the alignments with SAMtools for sorting, duplicate marking, and quality filtering (MAPQ ≥ 20). GATK HaplotypeCaller identified variants, including insertions, deletions, and single nucleotide variations (SNVs) using GATK’s hard-filtering parameters [26], followed by normalization with bcftools [27] and annotation with SnpEff and SnpSift [28]. Common variants across individuals or samples were identified using in-house Python scripts. Using a curated innate immune gene list from InnateDB (https://innatedb.com/)and a keyword-based search, we identified immune-related genes. We mapped the genes with variants to QTLs from Animal QTLdb (www.animalgenome.org), followed by functional enrichment of genes with variants using DAVID (https://david.ncifcrf.gov/home.jsp) v6.8 for GO Biological Processes and KEGG pathway enrichment and visualized for significant data (P-value < 0.05). Additional analysis such as runs of homozygosity (ROH) using PLINK [29] identified ROH islands that are genomic regions shared by >50 % of individuals or samples. Of the results obtained the genes in the top 1 % were considered. We detected selective sweeps using the Raised Accuracy in Sweep Detection tool (RAiSD) [30], and prioritized overlapping genes from ROH islands and RAiSD regions as candidate selection signatures.

### Data Collection and Quality Assessment

The whole-genome sequencing data for Nelore breed (n = 14) and Gir breed (n = 20) was retrieve from the NCBI Sequence Read Archive (SRA) using the SRA Toolkit (https://trace.ncbi.nlm.nih.gov/Traces/sra/sra.cgi?view=software). We assessed the quality of raw sequencing reads with FastQC [23], and filtered low-quality bases and adaptor contaminants using fastp [24] with the following parameters: cut front and cut tail to remove low-quality bases at read ends, cut mean quality 20 to trim when the average quality in a sliding window fell below 20, qualified quality phred20 and unqualified percent limit 30 to retain reads with ≥70 % of bases above Q20, length required 25 to discard reads shorter than 25 bp, n base limit 10 to remove reads containing more than 10 ambiguous bases, and detect adapter for pe to automatically detect and remove adapter sequences for paired-end data. We retained reads that passed quality thresholds based on Q20 and Q30 for downstream analyses, including alignment to the reference genome and variant calling, because these thresholds ensure base call accuracies of 99 % and 99.9 %, respectively, which minimizes sequencing errors and improves the reliability of downstream analyses.

### Read Mapping and Preprocessing

The high-quality sequence reads for the Nelore breeds and Gir breed to the *Bos taurus* reference genome (Hereford breed, ARS-UCD2.0) using BWA-MEM [25]. QUAST identified the assembly statistics for the reference genome. Using SAMtools [27], we processed the mapped reads to sort, correct mate information, mark duplicates, and index files. We assessed alignment quality with SAMtools stats and retained only reads with mapping quality (MAPQ ≥ 20) for variant calling, since this threshold corresponds to a 99% probability of correct mapping and ensures that aligned reads enter downstream analyses.

### Variant calling and Annotation

Three variants, including insertions, deletions and SNVs, were identified using GATK HaplotypeCaller [26]. The GATK’s recommended hard-filtering parameters were used for filtering the variants: for SNVs, Qual By Depth (QD) < 2.0, Fisher Strand (FS) > 60.0, RMS Mapping Quality (MQ) < 40.0, Strand Odds Ratio (SOR) > 3.0, Mapping Quality Rank Sum Test (MQRankSum) < – 12.5, and Read Pos Rank Sum Test (ReadPosRankSum) < –8.0; for Indels, QD < 2.0, FS > 200.0, SOR > 10.0, and ReadPosRankSum < –20.0 [31, 32]. Using bcftools norm, the variants were normalized and annotated them against the *Bos taurus* reference genome (ARS-UCD2.0, Ensembl GFF3 release 115) using SnpEff. Further, the genotype and gene-level information was extracted using SnpSift with vcfEffOnePerLine command. Variant calling and annotation were performed across the entire genome, including autosomes, sex chromosomes (X, Y), mitochondrial genome (MT), and unplaced scaffolds. Finally, the common variants across individuals or samples within each breed were identified using in-house Python scripts.

### Identification of Immune-Related Genes

To identify the immune-related genes with variations, we employed a combined approach using both curated database and keyword-based search, implemented through an in-house Python script. First, a curated list of innate immune genes (n = 1697) was obtained from InnateDB. However, since InnateDB focuses solely on innate immunity, we aimed to capture other immune-related genes that may contribute to adaptive immunity or immune regulatory functions but are not included in the database. To achieve this, we conducted a keyword-based search on the annotated gene list derived from the genome data. The following immune-related keywords were used for the search: immunoglobulin, immunoreceptor, autoimmune, Toll-like receptor (TLR), IgG, autoimmune, autophagy, immunogen, immune, innate, T-cell, B-cell, lymphocyte, histocompatibility, *CD24, CD4, LY96, IFIT3, PGLYRP1, NKG2D, UL16*, leukocyte, cytokine, antimicrobial peptide, beta-defensin 2, *IL15, IL2*, and chemokine. This combined approach allowed for a comprehensive identification of genes involved in immune-related functions.

### Analysis of Quantitative Trait Loci (QTL)

Using the genomic variations such as insertions, deletions, and SNVs, identified by GATK in both Nelore breed and Gir breed, we looked at the QTLs in the genes that had genomic variations. We used the QTL data of the Hereford breed (ARS_UCD2.0) in GFF format obtained from the cattle QTLdb of the Animal QTLdb database (www.animalgenome.org). The QTL deposited in the database were related to characters associated with reproduction, production of milk, health, exterior, and meat and carcass. The locus of a qualitative trait is shown as a 4-bp-long segment, referred to as a QTL span. Using an in-house Python script, we identified the reported QTLs associated with various traits in the genomic regions showing variants.

### Gene Ontology Enrichment Analysis

To investigate the functional relevance of genes with variants, we performed Gene Ontology (GO) enrichment analysis separately for genes impacted by insertions, deletions, and SNVs. The analysis was conducted using the DAVID Bioinformatics Resources (v6.8) [33]. For each variant category, a list of variant genes was submitted, and GO terms were evaluated under the categories of Biological Process (BP) and, KEGG Pathways. Only significantly enriched terms with a P-value < 0.05 were retained. These significantly enriched terms were visualized using Python’s matplotlib/seaborn package [34, 35].

### Runs of Homozygosity (ROH) Analysis

The runs of homozygosity (ROH) analysis in the Nelore breed (n = 14) and in the Gir breed (n = 20) were conducted using PLINK v1.9 [29]. We retained only autosomal SNVs, excluding sex chromosomes (X, Y), the mitochondrial genome (MT), and unplaced scaffolds, because these regions have different inheritance patterns, different recombination rates, and mapping reliability issues that could bias homozygosity estimates [36, 37]. The VCF files were converted to PLINK binary format and quality control was applied by filtering out SNVs with genotype call rate < 0.90 and minor allele frequency (MAF) < 0.05. We estimated ROH with a sliding-window approach of 50 SNVs using the following criteria: (i) at least 50 consecutive homozygous SNVs to define a ROH, (ii) minimum ROH length of 1 Mb, (iii) density of at least one SNV per 50 kb, (iv) maximum gap between consecutive SNVs of 1 Mb, and (v) allowance of up to one missing SNV and one heterozygous SNV per window. The ROH segments were summarized across individuals or samples within each breed. To identify the region of genome enriched for homozygosity called ROH islands, the individual ROH segments were converted to BED format and non-overlapping 50 kb windows was generated across the genome with bedtools [38]. Thus, ROH islands were defined as the overlap between ROH segments and windows and designated windows shared by more than 50% of individuals or samples. We annotated genes within these consensus ROH islands against the *Bos taurus* reference genome (Hereford breed, ARS-UCD2.0) and highlighted the top 1% of genes by overlap frequency as candidate genes under selection.

### Selective Sweep Identification using RAiSD (Raised Accuracy in Sweep Detection)

We conducted genome-wide scans for signatures of selection in Nelore and Gir breeds using the µ statistic implemented in RAiSD [30]. This composite measure integrates multiple signals of selection by jointly evaluating deviations in the site frequency spectrum (SFS), patterns of linkage disequilibrium (LD), and reductions in genetic diversity (VAR) across the genome. We extended genomic regions exhibiting strong selection signals by ±50 kb from each focal site. These regions were annotated using bedtools with the *Bos taurus* reference genome (Hereford breed, ARS-UCD2.0). Genes overlapping both RAiSD-identified regions and ROH islands were considered as putative candidate genes under selection.

## Results

### Reference Genome Statistics and Quality Assessment of Sequence Reads

The *Bos taurus* reference genome (Hereford breed, ARS-UCD2.0,) was used for this study. The *Bos taurus* reference genome consists of 1,958 contigs including 29 autosomes, X, Y, and Mitochondrial chromosomes, with a total length of 2.77 Gb, an N50 of 103.3 Mb, and minimal gap content (1.02 Ns per 100 kbp).

We analyzed whole-genome sequencing data comprising 14 Nelore and 20 Gir samples retrieved from the NCBI Sequence Read Archive. We assessed raw-read quality using Phred thresholds and filtered reads with fastp, retaining those with Q20 scores >95% and Q30 scores >85% (Fig. 2). After filtering, 96.74% of reads were retained and 3.26% were discarded. We mapped the filtered reads to the *Bos taurus* reference genome (Hereford breed, ARS-UCD2.0) using BWA-MEM and processed the alignments with SAMtools. The processed alignments showed alignment rates >95% and mapping quality (MAPQ) scores >50, confirming suitability for downstream variant discovery.

**Fig. 1.**
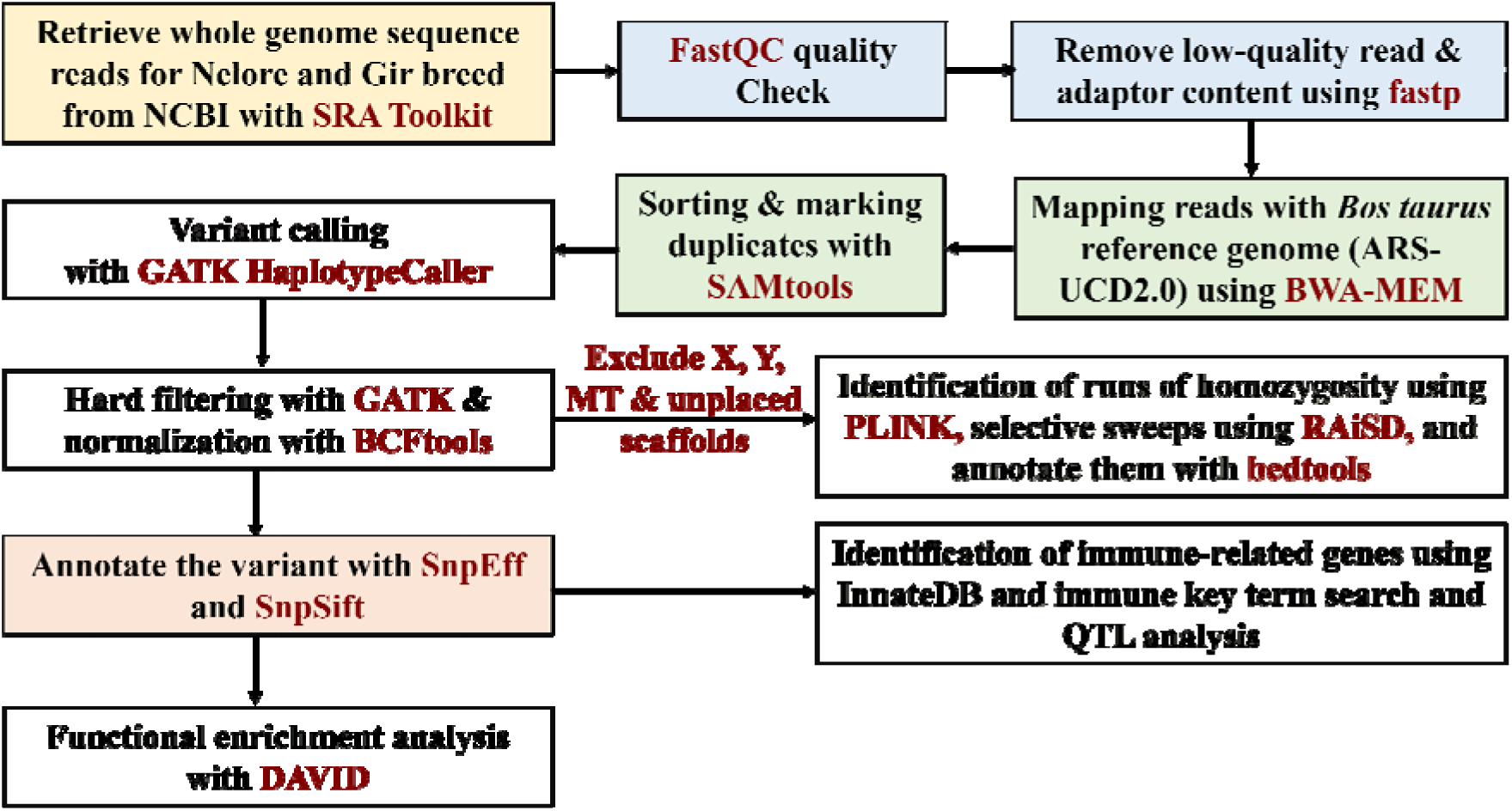
Workflow used in this study.

**Fig. 2.**
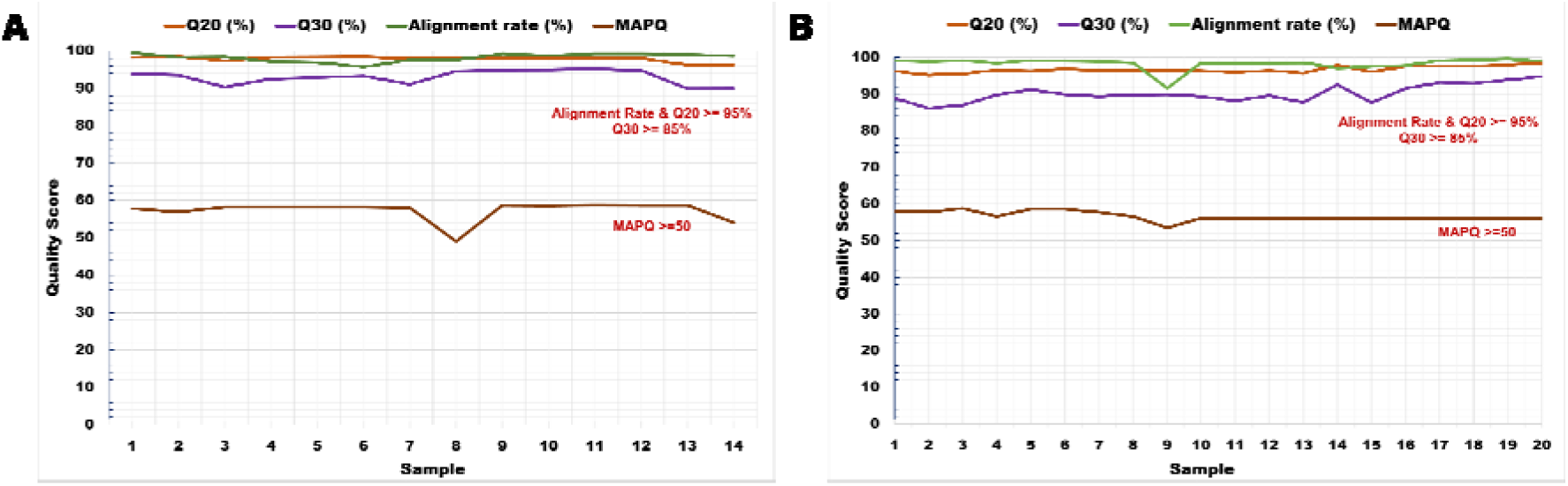
Quality assessment of whole-genome sequencing reads for Nelore (n = 14) and Gir (n = 20) breeds. (A) Phred quality scores (Q20 and Q30) after filtering with fastp, along with alignment rate and mapping quality (MAPQ) for the Nelore breed. (B) Phred quality scores (Q20 and Q30) after filtering with fastp, along with alignment rate and mapping quality (MAPQ) for the Gir breed.

### Variant Identification and Annotation

GATK identified 1,884,058 InDels (insertions and deletions) and 13,997,533 SNVs. The transition/transversion (Ti/Tv) ratio of SNVs was about 2.26 in the Nelore breed. Similarly, in the Gir breed, GATK identified 1,457,337 InDels (insertions and deletions) and 11,627,881 SNVs were detected (Fig. 3), with a Ti/Tv ratio of 2.25, which confirms the reliability of variant calling. Functional annotation showed that non-coding variants were abundant than coding variants, with intergenic variants (673,137 in Nelore breed; 121,610 in Gir breed) and intronic variants (269,240 in Nelore breed; 48,495 in Gir breed) as the most frequent categories. All variants also occurred in regulatory regions, including upstream, downstream, 3′ UTR, and 5′ UTR regions. Among variants in coding regions, missense variants numbered 1,437 in Nelore breed and 249 in Gir breed, followed by synonymous variants (1,535 in Nelore breed and 303 in Gir breed). High-impact alterations, though less frequent, included frameshift variants (37 insertions and 27 deletions in Nelore breed; 7 insertions and 3 deletions in Gir breed), stop-gained mutations (19 in Nelore breed; 2 in Gir breed), and splice-site variants (62 in Nelore breed; 8 in Gir breed) (Table 1). These missense, frameshift, and splice-site variants are likely to influence gene function.

**Fig. 3.**
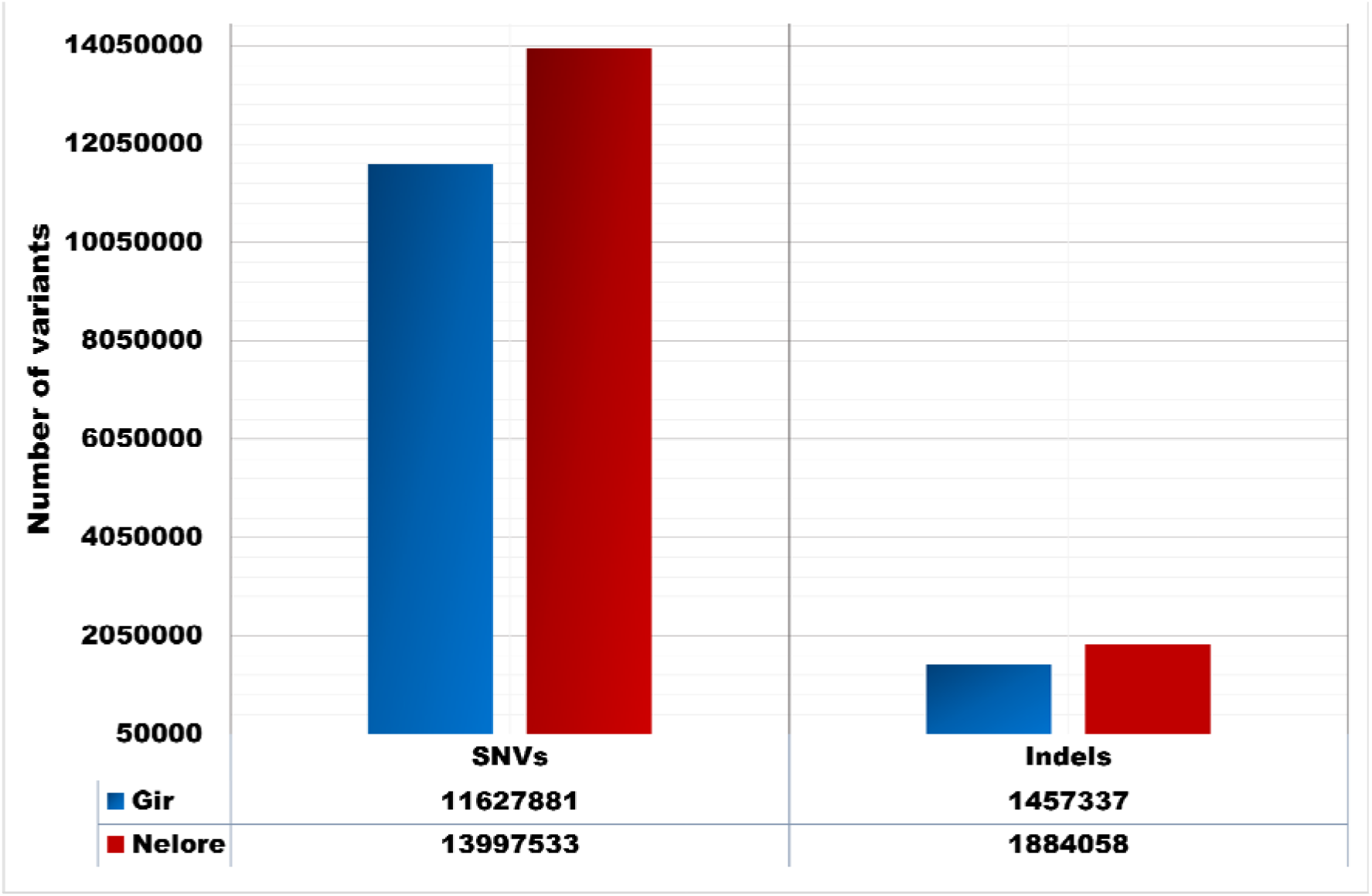
Number of variants, including InDels (insertions and deletions) and SNVs, identified in Nelore and Gir breeds using GATK after applying hard filtering parameters. The red bars represent variants in the Nelore breed, while the blue bars represent variants in the Gir breed.

**Table 1:** Functional classification of variants in Nelore and Gir breeds showing the number of genes annotated in different genomic regions as identified by snpEff. The annotations span different genomic regions, including non-coding regions, coding regions, regulatory elements, splice sites, and untranslated regions.

| Effects | Nelore SNV | Nelore Insertion | Nelore Deletion | Gir SNV | Gir Insertion | Gir Deletion |
| --- | --- | --- | --- | --- | --- | --- |
| Missense variant | 1437 | 0 | 0 | 249 | 0 | 0 |
| Frameshift variant | 0 | 37 | 27 | 0 | 7 | 3 |
| Start lost | 1 | 0 | 1 | 3 | 0 | 0 |
| Stop lost | 17 | 0 | 0 | 1 | 0 | 0 |
| Stop gained | 19 | 1 | 0 | 2 | 0 | 0 |
| Stop retained variant | 1 | 0 | 0 | 1 | 0 | 0 |
| Transcript ablation | 0 | 0 | 1 | 0 | 0 | 0 |
| Splice region variant | 426 | 43 | 35 | 63 | 10 | 5 |
| Splice acceptor variant | 11 | 7 | 6 | 5 | 73 | 1 |
| Splice donor variant | 13 | 9 | 6 | 2 | 2 | 1 |
| Intergenic region | 673137 | 11048 | 8754 | 121610 | 4403 | 2805 |
| Intron variant | 269240 | 6348 | 4641 | 48495 | 2228 | 1424 |
| Downstream gene variant | 40445 | 2613 | 1645 | 7011 | 507 | 263 |
| Upstream gene variant | 41672 | 0 | 1704 | 6993 | 498 | 252 |
| Intragenic variant | 4658 | 223 | 151 | 977 | 53 | 45 |
| 3_prime UTR variant | 6000 | 486 | 296 | 1121 | 101 | 54 |
| 5_prime UTR variant | 3698 | 203 | 127 | 633 | 41 | 26 |
| 5_prime UTR premature start codon gain variant | 625 | 0 | 0 | 94 | 0 | 0 |
| Non coding transcript exon variant | 5624 | 307 | 201 | 1133 | 73 | 39 |
| Non coding transcript variant | 0 | 0 | 1 | 0 | 0 | 0 |
| Synonymous variant | 1535 | 0 | 0 | 303 | 0 | 0 |
| Disruptive inframe insertion | 0 | 8 | 0 | 0 | 3 | 0 |
| Conservative inframe insertion | 0 | 1 | 0 | 0 | 1 | 0 |
| Conservative inframe deletion | 0 | 0 | 1 | 0 | 0 | 1 |
| Disruptive inframe deletion | 0 | 0 | 5 | 0 | 0 | 0 |

We observed that variants, including insertions, deletions, and SNVs, were distributed across autosomes, sex chromosomes (X and Y), the mitochondrial genome, and unplaced scaffolds in both Nelore and Gir breeds (Fig. 4). In the Nelore breed, genes with SNVs and insertions were most abundant on chromosome 3, with 1,683 genes (Fig. 4A) and 876 genes (Fig. 4B), respectively, while deletions were most frequent on chromosome 5, with 702 genes (Fig. 4C). In the Gir breed, genes with SNVs were most frequent on chromosome 5, with 1,156 genes (Fig. 4D), whereas insertions and deletions were most common on chromosome 1, with 453 genes (Fig. 4E) and 288 genes (Fig. 4F), respectively.

**Fig. 4.**
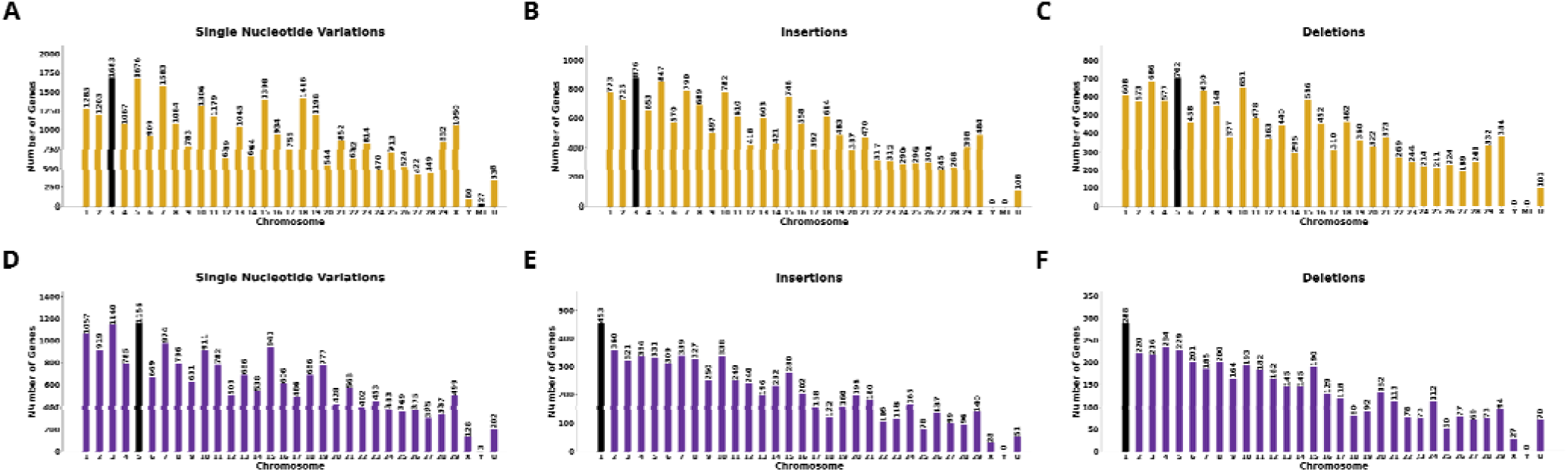
Distribution of variants associated genes across autosomes, sex chromosomes (X and Y), the mitochondrial genome, and unplaced scaffolds in Nelore and Gir breeds. Panels A–C show variant distributions in Nelore breed: (A) Genes with single nucleotide variants, (B) Genes with insertions, and (C) Genes with deletions. Panels D–F show variant distributions in Gir breed: (D) Genes with single nucleotide variants, (E) Genes with insertions, and (F) Genes with deletions. Black bars indicate the maximum variant counts.

We found that the Nelore breed harbored a relatively large number of variant-affected genes than the Gir breed. Because our study focused on immune-related genes, we applied two identification strategies: we retrieved innate immune genes from the InnateDB database and we identified additional immune genes not listed in InnateDB by keyword-based searches. In the Nelore breed (Table 2), 26,953 genes carried SNVs, including 1,083 immune-related genes, of which 719 were innate immune genes. Insertions affected 15,622 genes, including 603 immune-related genes (402 innate), and deletions occurred in 12,385 genes, including 512 immune-related genes (339 innate). In the Gir breed (Table 2), 19,117 genes carried SNVs, including 759 immune-related genes (502 innate). Insertions occurred in 6,427 genes, including 220 immune-related genes (154 innate), and deletions occurred in 4,216 genes, including 145 immune-related genes (98 innate). These results indicate that the Nelore breed carries a larger burden of variants in immune-related genes than the Gir breed, suggesting differences in immune gene evolution between the two breeds.

**Table 2.** Number of genes, including immune-related genes (innate immune genes identified using InnateDB), carrying variants in Nelore and Gir breeds.

| Breed | Variants | Total genes | Immune genes | Innate genes |
| --- | --- | --- | --- | --- |
| Nelore | SNVs | 26953 | 1083 | 719 |
| Nelore | Insertions | 15622 | 603 | 402 |
| Nelore | Deletions | 12385 | 512 | 339 |
| Gir | SNVs | 19117 | 759 | 502 |
| Gir | Insertions | 6427 | 220 | 154 |
| Gir | Deletions | 4216 | 145 | 98 |

#### Variations involving insertions

SnpEff classified the variants into four categories: High impact, Moderate impact, Modifier impact, and Low impact. High-impact variants cause protein truncation; moderate-impact variants alter protein function; modifier variants occur in non-coding or regulatory regions (including UTRs); and low-impact variants have neutral effects. We identified 2,618 high-impact, 103,309 moderate-impact, 37,416,134 modifier-impact, and 237,935 low-impact variants in the Nelore breed. In the Gir breed we observed 2,244 high-impact, 86,542 moderate-impact, 31,005,968 modifier-impact, and 196,473 low-impact variants. These counts represent the distribution of variant impact classes across the genome. For downstream analysis, we focused only on genes that had high, moderate, and modifier variants.

We found 45 genes with high-impact variants in the Nelore breed, including immune-related genes. The immune-related genes included Toll-like receptor 3 (*TLR3*) and LOC508441 (*CD33*), a member of the immunoglobulin-like domain family, both affected by a frameshift variant. We also identified high-impact insertions in non-immune genes such as *METTL6, VWDE, EP300, KIF13B, ZNF484, TTLL5, MYCBP2, PHKB, BDP1, OSMR, CDH18*, and *EFCAB13*. Fifteen genes had moderate-impact variants consisting of both disruptive and conservative inframe insertions. Non-immune genes with disruptive inframe insertions included *ABCA13, ZNF280D, UNC13C, OR5M5, BRCA1, ARL15*, and LOC529488. Furthermore, 15,609 genes harbored modifier variants, of which 602 were immune-related, including *RCAN1, IFNAR1, TP63, IL1RAP, IFI6, TRIM42/63, BCL10, IFNG*, and *IRAK3*.

Variations more than 50 base pairs (bp) can significantly affect gene expression and function by altering gene dosage, disrupting coding regions, or interfering with long-range regulation [39], we focused on genes impacted by variants exceeding 50bps long. A total of 37 genes had with insertions longer than 50 bp (Fig. 5). The gene LOC112444661 on chromosome 27 exhibited the largest length of variation (LOV) at 76 bp, followed by LOC781770 on chromosome 24 (74 bp), LOC112444213 on chromosome 24 (74 bp), and LOC132345508 on chromosome 6 (72 bp). In addition, several other genes showed insertions more than 50 bp, including *GRXCR1* (72 bp, chromosome 6), *DNAH7* (66bp, chromosome 2), *TMEM232* (64 bp, chromosome 7), *ZNF277* (63 bp, chromosome 4), *TBX15* (62 bp, chromosome 3), *HDAC9* (57 bp, chromosome 4), *MAGI2* (55 bp, chromosome 4), and *FOXP2* (55 bp, chromosome 4). Among immune-related genes, *DOCK4* on chromosome 4 had an insertion of 63 bp. Furthermore, several innate immune genes exhibited insertions longer than 20 bp, including *PRKDC, CXCR4, CDK6, IGF2, PIK3R1, ANKRD17, CDKN2A, IGF1R, SYK, NLRX1, TCF4, TLR2, RCOR1, TRIM38, IL1RAP, FANCC, MUL1, MAP3K4/MAP3K14, OPTN*, and *IRF2*. In addition, some genes had multiple independent insertions at different site. These included *TP63*, which contained two insertions of 10 bp each, and *DDX1* had two insertions of 12 bp and 11 bp. The gene *PIK3R1* had three independent insertions, one of 36 bp and two of 16 bp each.

**Fig. 5.**
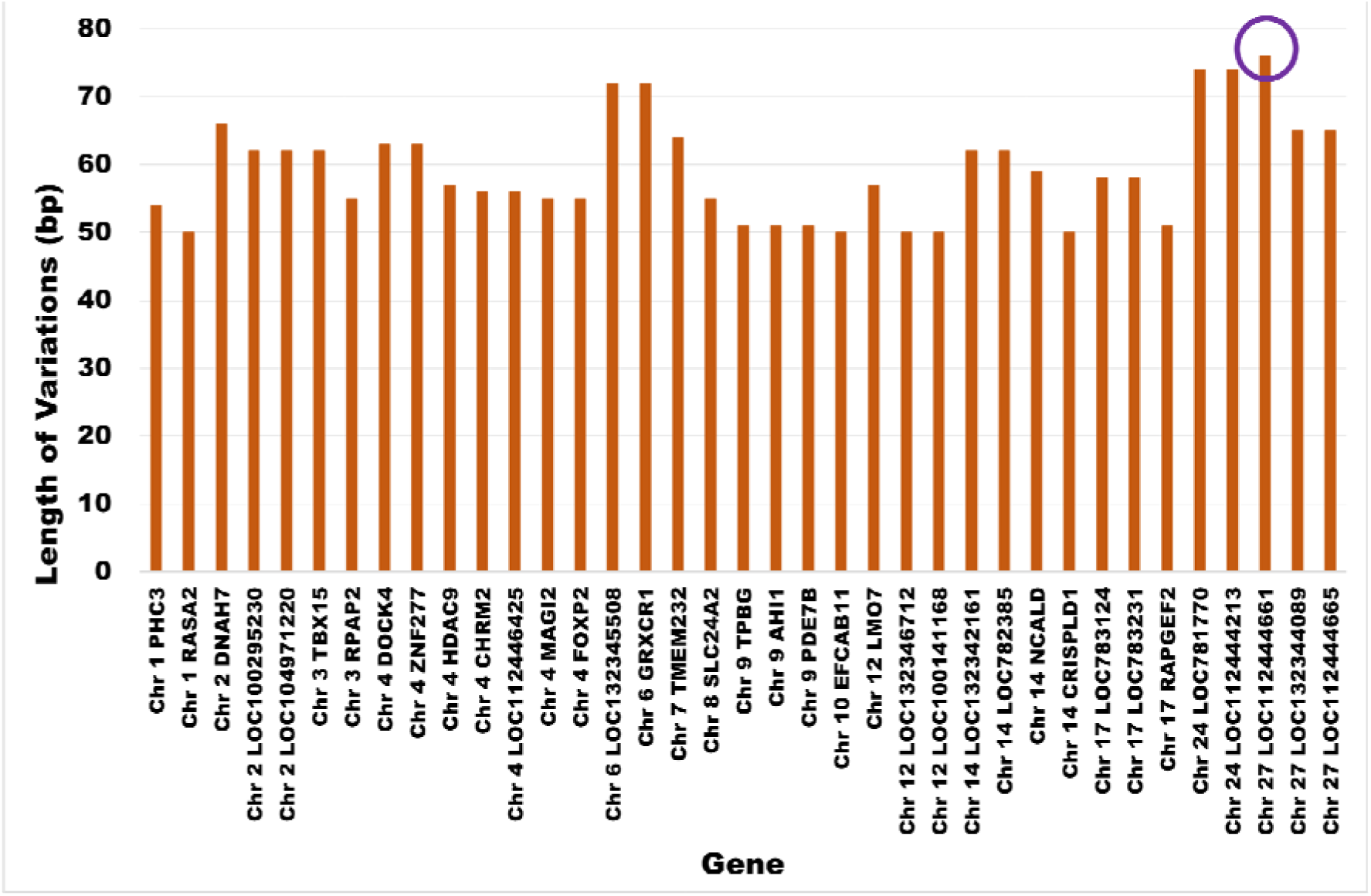
Characters of 37 genes with insertions longer than 50 bp in Nelore breed. The circle marks the gene with highest number of insertions

We identified nine genes with high-impact frameshift variants in the Gir breed, including *VWDE, OR6C63, CLEC12A, GPRC6A, ZNF432, OSMR*, and *H4C19*. The innate immune gene *TLR3* and the immune-related gene LOC508441 (identified by keyword search) also carried high-impact frameshift variants, as observed in the Nelore breed. Four non-immune genes carried moderate-impact variants; disruptive inframe insertions occurred in *ZNF280D, OR52Z19*, and LOC104976340 (located on an unplaced scaffold). A total of 6,416 genes carried modifier variants, of which 219 were immune-related, including *JAM3, NOX4, ETS1, IFIT5, PTEN, BCL2, TCF4, PIK3C3, TINAG, TRAF3, NFZT5, DOCK9, PCBP1, TLR4, JAK2, IRAK3, TAFA2, TP63, ALCAM, PROS1*, and *RCAN1*. Consistent with the Nelore breed, we observed genes with length of variation (LOV) more than 50 bp in the Gir breed. Three genes had such long variants: LOC132345096 on chromosome 4 (77 bp), LOC112444661 on chromosome 27 (76 bp), and *DNAI7* on chromosome 5 (52 bp). Among immune-related genes, 11 carried longer variants, including *ADCY8, RNF125, PLXNA4, IL1A, SOCS2, TINAG, TAFA2, MSR1, TRIL, PARD3*, and *MUL1*.

Overall, we identified 5,542 genes with insertions shared between Nelore and Gir breeds, while 10,080 genes were unique to Nelore breed and 885 genes were unique to Gir breed. Four hundred twelve immune-related genes, including *IFNAR1, CD47, GSK3B, FXR1, PTX3, RFTN1, PIK3CB, CXCR4, MARCO*, and *BCL10*, were specific to Nelore breed, and 29 immune-related genes, including *TYRO3, NFATC4, MR1, C1QBP, RANBP9, SOCS6, IFIT5*, and *IKBKB*, were specific to Gir breed.

#### Variations involving deletions

In the Nelore breed, we identified 12,385 genes had deletions, including 35 with high-impact variants, 6 with moderate-impact variants, and 12,371 with modifier variants. High-impact frameshift deletions were observed in genes such as *HACL1, DNAH11, DPY19L2, ZNF804B, CC2D2A, MTFR2, OR4C27, VPS16, TOX, OR5M5, PER3, OR5F2B, KRT34, DCTN5, PDZD8, TOP6BL*, and the innate immune gene *JAM3*. Moderate-impact disruptive in-frame deletions were present in non-immune gene such as *N4BP2L2, OR4C27, BDP1*, and LOC783920. Deletion events also affected the regulatory and non-coding regions of 512 immune-related genes, including 339 innate immune genes such as *IFNAR1, PROS1, TRAT1, CD200, GSK3B, IL1RAP, TP63* and others. We further observed 122 genes with deletions longer than 50 bp (as shown in Fig. 6), with LOC101902469 on chromosome 18 showing the longest deletion (136 bp; splice-site variant, high impact) and LOC112449180 on chromosome 12 had a 98 bp deletion (annotated as transcript ablation, a high impact variant). Long deletions were also found in regulatory regions of immune and non-immune genes. The immune-related genes including *PROS1* (66 bp, chromosome 1), *TNIP3* (52 bp, chromosome 17), *NLRP12* (46 bp, chromosome 18), and *DOCK4* (40 bp, chromosome 4). Multiple intronic deletions were detected in *TNIP3* and *DOCK4*.

**Fig. 6.**
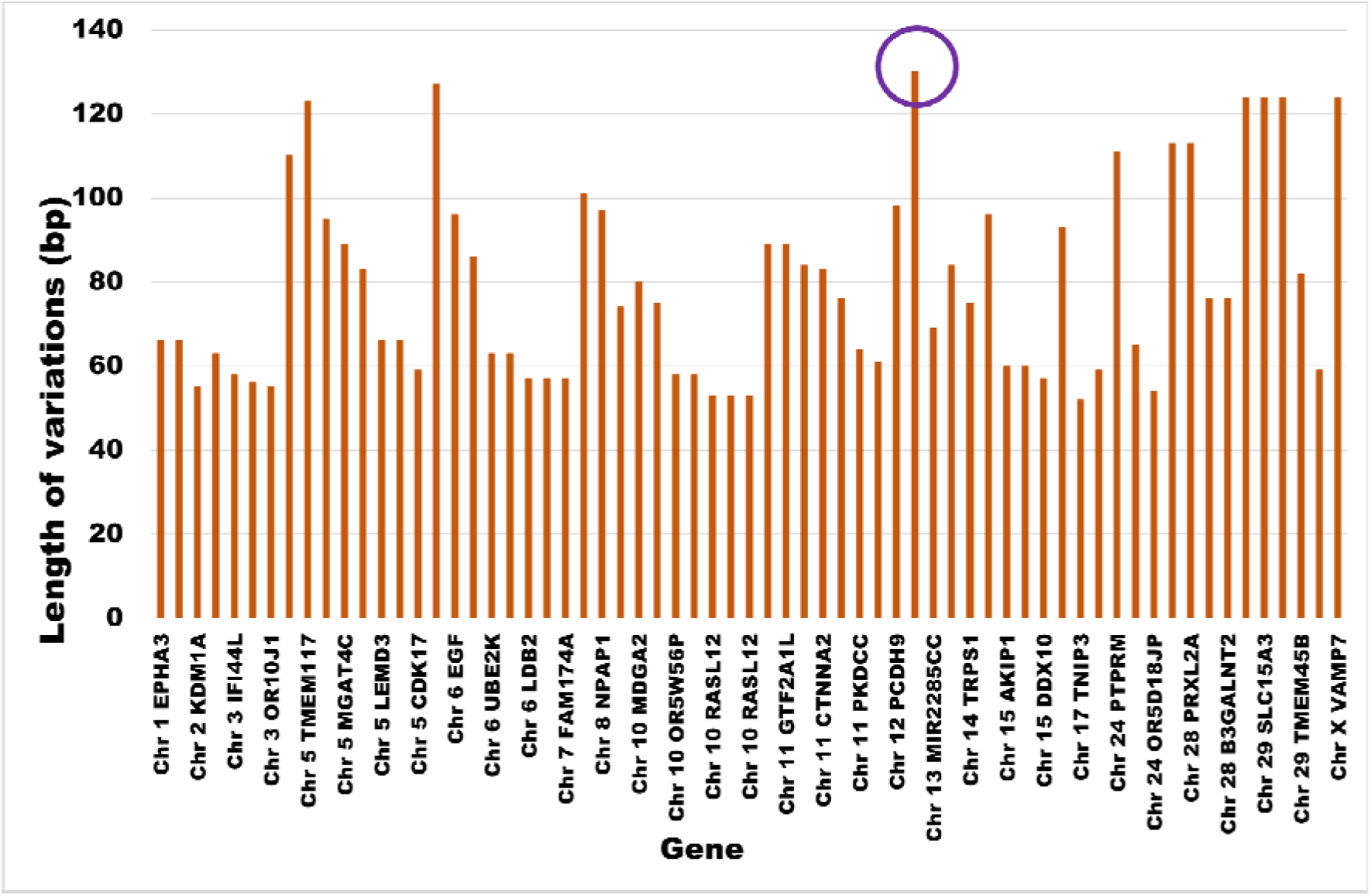
Variations of 122 genes with deletions longer than 50 bp in Nelore breed. The circle marks the gene with highest number of deletions.

In the Gir breed, we identified 4,216 genes with deletions, including 4 high-impact variants, 1 moderate-impact variant, and 4,213 modifier variants. High-impact deletions included splice-site variants in *CNPPD1* and frameshift variants in *DNAH11, SARNP*, and LOC107132987. The immune-related gene LOC100138951 (identified by keyword search) corresponding to *PAX5*, had frameshift variant. *PAX5* is a B-cell specific transcription factor involved in the adaptive immune system [40]. The moderate-impact conservative in-frame deletion in LOC100138951. No immune-related genes had moderate-impact deletions was found in Gir breed. Deletions were also observed in regulatory and non-coding regions of 144 immune-related genes, including 98 innate immune genes such as *PROS1, GSK3B, TP63, MARCO, JAM3*, and others. Furthermore, 12 genes had deletions longer than 50 bp, such as *ELP4* (107 bp, chromosome 15), *LEMD3* (66 bp, chromosome 5), *FAT3* (59 bp, chromosome 29), *EXD1* (54 bp, chromosome 10), *ELK3* (51 bp, chromosome 5), *GALNT18* (51 bp, chromosome 15), and LOC132342331 (51 bp, chromosome 15). Immune-related long deletions included *PROS1* (42 bp, chromosome 1) and *SOCS2* (18 bp, chromosome 5).

Overall, we observed 3,417 genes with deletions shared between Nelore and Gir breeds. Additionally, 8968 genes were specific to Nelore breed and 799 genes were specific to Gir breed. Among these, 390 immune-related genes were unique to Nelore breed, including *IFNAR1, ALCAM, CD47, ATG3, MASP1, ADIPOQ, NOD1, IRAK3, TLR4/10, FOXA2, HSPA14*, and *NFATC2* and 23 immune-related genes were specific to Gir breed, such as *DHX36, BCL2, MAPK1, AP3B1, CXCL9, IL22RA1, F2RL1, IGF1*, and *IL12B*.

#### Variations involving single nucleotide variations

We identified 26,953 genes with SNVs in the Nelore breed, comprising 64 high-impact variants, 986 moderate-impact variants, and 26,925 modifier variants. High-impact variants, including splice-site mutations and variants in start and stop codons, occurred in genes such as *LRRC31, TRPC1, ZNF445, MEG8, DNAH12*, and the immune-related genes *CD46* and *IL26*. Moderate-impact missense variants occurred in 916 non-immune genes and 45 immune-related genes, including innate immune genes such as *NLRP12, MAP3K1, CD46, IL1RL2, IFNGR1, ANXA1, IFI6, ITGAM, MAPK8, TECPR1, ELP2*, and *NCKAP1L*. Many SNVs occurred in non-coding and regulatory regions, affecting 25,846 non-immune genes and 1,079 immune-related genes. We observed 120 genes with more than 500 SNVs each. Among them, *LRRC7* (1,679 variants in UTRs and regulatory/non-coding regions) showed the highest SNV frequency. Other examples include *UNC13C* (849; stop lost and missense variants), *DNM3* (813; stop lost), and genes with missense variants *DPYD* (512), *BEND5* (654), *MAGI2* (682), *RELN* (899), *BBS*9 (739), *SOX5* (649), *VPS13A* (815), and *ADAMTSL1* (668). Genes with regulatory-region SNVs included *MACROD2* (1,075), *TCF12* (1,072), *TRNAK* (937), *FOXP2* (915), *LINGO2* (907), *PARD3B* (606), *CDH18* (531), *HDAC9* (537), *MICU1* (504), and *NRG3* (546).

We identified 19,117 genes with SNVs in the Gir breed, comprising 6 high-impact variants, 220 moderate-impact variants, and 19,088 modifier variants. High-impact variants occurred in genes such as *PIK3R3, ZNF217, CC2D2B, CCDC73, OR6C7G*, and the innate immune gene *PLG*. Moderate-impact missense variants occurred in 195 non-immune genes (for example, *AOX1, OR5H13, TFRC, DLG1, FCRL3*) and in five innate immune genes, including *DHX36, MARCO, IFNGR1, AKAP10*, and *DEFB7*. SNVs also occurred in the regulatory and non-coding regions of 17,511 non-immune genes and 759 immune genes. Unlike the Nelore breed, no genes had more than 500 SNVs in the Gir breed.

We observed that 17,469 genes with SNVs were shared between Nelore and Gir breeds. We found 9,486 genes specific to the Nelore breed, including 382 immune-related genes such as *IFNAR1, NRIP1, NFKBIZ, ATG3, ADIPOQ, SENP2, IL12A*, and *IFIH1*. In contrast, we identified 1,648 genes specific to the Gir breed, including 58 immune-related genes such as *IFNGR2, SIAH2, ACKR4, MX2, MIRLET7C, MTOR, BIRC5, TRIM65, TIRAP, JAM3, TOLLIP, HRAS*, and *IFITM1*.

### Multi-Variant Genes in Nelore and Gir breeds

A total of 9327 genes had all three types of variants, namely insertions, deletions, and SNVs in Nelore breed. Among them, 364 were immune-related genes, of which 253 genes were classified as innate immune genes based on the InnateDB database. These included key immune genes such as *IFNAR1, PROS1, ALCAM, CD47, IL1RAP, MASP1, ADIPOQ, TRIM42, PIK3CB, STAG1,* and *MARCO*. *TLR3* gene exhibited insertions resulting in frameshift and splice-site variants, deletions in non-coding intronic regions, and SNVs distributed across regulatory regions (upstream, downstream, and intronic). Similarly, CD46 had a stop-gained SNV, while insertions and deletions were distributed across its regulatory region. Missense SNVs were observed in several immune genes including *IFI6, CTSS, NCKAP1L, ANXA1, MAP3K5/1, IFNGR1, IL1RL2, C4BPB, CD46, NLRP12, CTSH, TREML2,* and *ITGAM*, whereas the insertions and deletions in these genes were present in non-coding regulatory regions.

The presence of multiple variant types within the same gene is particularly important, as it suggests cumulative or synergistic effects on gene regulation and protein function [41]. In the Gir breed, 2552 genes had all three types of variations, including 76 immune-related genes, of which 47 were innate immune genes. In the Gir breed, SNVs, insertions, and deletions were commonly observed in the regulatory region of immune genes such as *PROS1, TP63, NR3C1, EDIL3, JAK2, MAP3K1, PIK3C3, TCF4, BCL2, THRB, NOX4*, and *JAM3*.

### Mapping of quantitative trait loci (QTL)

We mapped QTLs to genes carrying insertions, deletions, and SNVs identified with GATK in Nelore and Gir breeds. For insertions, we mapped QTLs for 5420 genes in the Nelore breed, including 284 immune genes, of which 203 are innate immune genes. These genes associated with multiple QTL categories: health (1295 genes), reproduction (2061 genes), milk (3196 genes), meat and carcass (1960 genes), production (1821 genes), and exterior (975 genes). Among the immune genes, 81 were linked to health traits, 169 to milk, 100 to meat and carcass, 95 to production, 108 to reproduction, and 61 to exterior traits. Several genes were mapped to multiple QTLs, with variants distributed across coding, non-coding, and regulatory regions. We focused primarily on coding variants, such as high- and moderate-impact variants, because they are likely to affect biological function and protein activity. In the Nelore breed, the immune gene LOC508441 carried a high-impact variant and linked to production and reproduction QTLs, while no QTL mapped to *TLR3*. Non-immune genes included *METTL6* linked to health, *VWDE* linked to meat and carcass, and *EP300* linked to health and production. Notably, *EP300* and *PHKB* linked to health, meat, and carcass, and *OSMR* linked to health, meat, milk, and production harbored frameshift and splice-site mutations, suggesting pleiotropic effects across multiple traits. Genes such as *KIF13B, GPCPD1*, and *TMEM132B* mapped specifically to milk-related QTLs, while *MYCBP2, TTLL5*, and *CDH18* associated with carcass and production traits. BDP1 showed multiple splice variants linked to exterior, health, and reproduction, whereas *UNC13C* and *BRCA1* harbored disruptive inframe insertions associated with carcass traits.

In the Gir breed, we mapped QTLs for 2,425 genes, including 124 immune genes, of which 92 are innate. These genes distributed across health (674 genes), reproduction (971 genes), milk (1511 genes), meat and carcass (1071 genes), production (972 genes), and exterior (553 genes). Among immune genes, 39 linked to health, 96 to milk, 50 to meat and carcass, 52 to reproduction, 61 to production, and 36 to exterior traits. Several genes had high- and moderate-impact variants overlapping key QTLs. Non-immune *VWDE* harbored a frameshift variant associated with carcass traits, LOC508441 carried a frameshift variant linked to both production and reproduction, and *ZNF280D* had a disruptive inframe insertion tied to exterior, health, and production. *OSMR* exhibited multiple high-impact splice variants and intronic mutations that overlapped QTLs for health, carcass, milk, and production, underscoring its pleiotropic role.

We mapped QTLs for deletions to 4,380 genes in the Nelore breed, including 239 immune genes, of which 171 are innate immune genes. These genes associated with QTL categories as follows: health (1,111 genes), reproduction (1,729 genes), milk (2,604 genes), meat and carcass (1,658 genes), production (1,522 genes), and exterior (826 genes). Among the immune genes, 64 linked to health traits, 142 to milk, 88 to meat and carcass, 88 to reproduction, 76 to production, and 70 to exterior traits. Several genes carried high- and moderate-impact variants that overlapped QTLs. No QTLs were mapped to the innate immune genes. *JAM3* had a frameshift variant. Non-immune genes such as *DNAH11, ZNF804B, TOX,* and LOC101902469 carried multiple splice and frameshift variants linked to health, milk, production, reproduction, and carcass traits, indicating pleiotropic effects. *CC2D2A and KRT34* associated specifically with carcass and milk traits, respectively. *BDP1* had a disruptive inframe deletion linked to exterior, health, carcass, production, and reproduction.

In the Gir breed, we mapped QTLs for 1,593 genes, including 77 immune genes, of which 54 are innate immune genes. These genes associated with QTL categories as follows: health (478 genes), reproduction (656 genes), milk (998 genes), meat and carcass (744 genes), production (655 genes), and exterior (362 genes). Among the immune genes, 25 were linked to health, 50 to milk, 30 to meat and carcass, 29 to reproduction, 37 to production, and 25 to exterior traits. No QTLs were mapped to the immune-related gene LOC107132987 (*PAX5*). Non-immune genes such as *DNAH11* had splice and frameshift variants linked to carcass and reproduction, and SARNP linked to reproduction traits.

For SNVs, we mapped QTLs to 77783 genes in the Nelore breed, including 464 immune genes, of which 333 are innate immune genes. These genes associated with health (1,753 genes), reproduction (2,535 genes), milk (4,676 genes), meat and carcass (2,551 genes), and exterior (1,266 genes). Among the immune genes, 127 were mapped to health traits, 277 to milk, 142 to meat and carcass, 145 to reproduction, and 82 to exterior traits. We identified 429 genes with high- and moderate-impact variants, including several innate immune genes. The innate immune gene *CD46* had a high-impact stop-gained variant and multiple missense variants, and mapped to QTLs for health, carcass, and reproduction, indicating a pleiotropic role. Among immune-related genes identified by keyword search, *IL26* had a splice donor variant linked to reproduction, and LOC100849046 had a splice acceptor variant linked to production and exterior traits. *IFI6, CTSS, NCKAP1L, MAP3K5, IL6ST, C7*, and *TECPR1* harbored missense variants associated with health, milk, reproduction, and production traits.

In the Gir breed, we mapped QTLs to 6,192 genes, including 343 immune genes, of which 253 are innate immune genes. These genes associated with health (3,753 genes), reproduction (2,112 genes), milk (3,753 genes), meat and carcass (2,252 genes), production (2,029 genes), and exterior (1,093 genes). Among the immune genes, 97 were linked to health, 205 to milk, 116 to meat and carcass, 117 to reproduction, 106 to production, and 68 to exterior traits. We identified 106 genes with high- and moderate-impact variants, including several innate immune genes. No QTLs were mapped to high-impact variants in innate immune genes such as *PLG* (start lost) and LOC528132 (stop gained). The innate immune gene *DHX36* had missense variants associated with exterior and production traits; *MARCO* associated with meat and carcass traits; and *IFNGR1* associated with milk traits. No QTLs mapped to innate immune genes *DEFB7* and *AKAP10*. Immune-related genes identified by keyword search included LOC100849046, linked to exterior and production; *IL36G*, linked to milk; and *IL36B*, linked to exterior, health, milk, production, and reproduction.

### Runs of Homozygosity (ROH) Analysis

We identified runs of homozygosity (ROH) islands in the Nelore breed as genomic regions shared by more than 50% of individuals or samples. The top 1% of these ROH islands encompassed 15 genes. A region shared by nine individuals or samples contained *ANKRD11, RNF166*, and *SPG7*, which are associated with milk traits; *CPNE7, DPEP1*, and *ZNF469*, which link to milk production; *ACSF3*, which associates with meat, carcass, and reproduction traits; *CDH15*, which associates with milk and reproduction; and *BANP*, which links to health. The innate immune genes *ZFPM1* (linked to health) and *CYBA* also occurred within ROH islands. Regions shared by eight individuals or samples included *MAGI2* (linked to meat, carcass, and reproduction), *GNAI1, MC1R, TCF25* (linked to health), *FANCA* (linked to milk and reproduction), and *DEF8* (linked to meat and carcass).

In the Gir breed, ROH analysis identified ROH islands comprising 141 genes. All 20 individuals or samples shared a homozygous region containing *ORIF1* (linked to milk), *ZNF262* (linked to exterior, milk, and production), and *ZNF200* (linked to milk). The innate immune gene *MEFV*, associated with milk, exterior, and production traits, also occurred within this shared region.

### Selective Sweep Detection

Genome-wide scans revealed multiple peaks of the µ statistic across all autosomes in Nelore and Gir breeds (Fig. 7). Manhattan plots show distinct peaks corresponding to genomic regions with strong signals of positive selection in each breed.

**Fig. 7.**
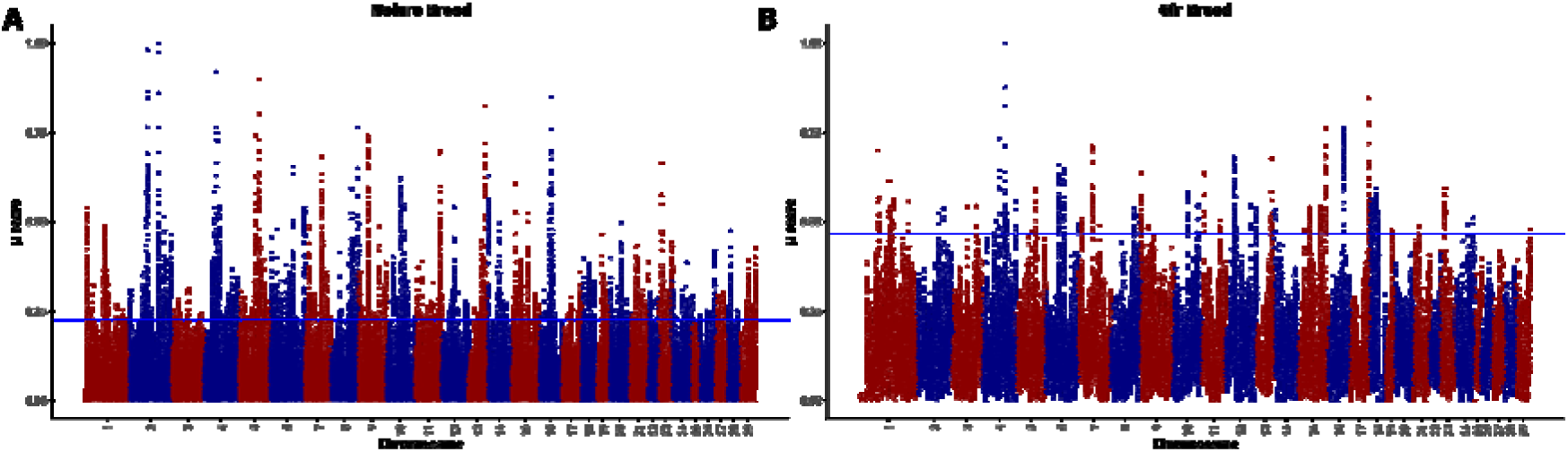
Genome-wide selective sweep detection using RAiSD. Genome-wide scans of the µ statistic across all autosomes in Nelore (A) and Gir (B) breeds. The horizontal blue line indicates the top 1 % threshold used to define candidate selective sweep regions.

We applied a significance cutoff corresponding to the top 1% of µ values to define candidate regions (horizontal line in Fig. 7). In the Nelore breed (Fig. 7A), numerous tall, narrow, and scattered peaks exceeded this threshold, indicating multiple strong, localized signals of positive selection. In the Gir breed (Fig. 7B), fewer peaks rose above the cutoff and they appeared less clustered. Peaks above the threshold in Nelore breed yielded 707 candidate genes, including 33 innate immune genes such as *CXCR1, IFI6, ARHGAP15, IFRD1, FOXO3, TRIM9, ATG5, PLXNA4*, and *CNOT4*. In Gir breed,

the top 1% cutoff highlighted 165 candidate genes, of which seven were innate immune genes, including *TP73, CSF2RB, NFAT5, NFATC3, PLCG2, COLEC12*, and *TAX1BP1*; the two additional immune-related genes identified by keyword search were *IK* and *PTCRA*. The sharp, high-amplitude peaks above the cutoff represent putative selective sweeps, whereas the broader signal distribution reflects background variation. Overall, both breeds harbor genomic regions under strong selection that are enriched for immune-related genes and loci linked to milk, production, reproduction, and health traits, but selection in the Nelore breed affects genomic regions and is sharply defined than in the Gir breed.

### Overlap of ROH Islands and Selective Sweeps

To identify strong candidate genes under selection, we assessed overlaps between the top 1% ROH islands and RAiSD-identified regions. In the Nelore breed, six genes overlapped both categories: *ANKRD11* (no QTL mapped), *MAGI2* (linked to meat, carcass, milk, production, and reproduction), LOC132345096 (linked to milk), *FOXP2* (linked to meat, carcass, and reproduction), *TCF12* (linked to milk, production, and reproduction), and *ATP5PO* (linked to meat and carcass). In the Gir breed, the genes *MEFV* (linked to milk, production, and exterior) and *ORIF1* (linked to milk) overlapped both ROH islands and RAiSD-identified regions. These overlapping genes represent strong candidates for selection signatures in the Nelore and Gir breeds.

### Gene Ontology (GO) and Pathway Enrichment analysis

We performed GO term enrichment and KEGG pathway analysis with DAVID for all genes carrying high-impact, moderate-impact, and modifier-impact variants (insertions, deletions, and SNVs) in Nelore and Gir breeds. We considered terms with P-value < 0.05 as significant.

For genes with insertions in the Nelore breed, enriched biological processes included regulation of transcription by RNA polymerase II, signal transduction, intracellular signal transduction, protein phosphorylation, and DNA repair (Fig. 8A). Immune-related GO terms were also enriched, including T cell receptor signaling, negative regulation of NF-kappaB transcription factor activity, inflammatory response, and endocytosis. KEGG pathway enrichment (Fig. 8B) highlighted neuroactive ligand-receptor interaction, PI3K-Akt signaling, JAK-STAT signaling, Rap1 signaling, and cAMP signaling. Immune-related pathways such as MAPK signaling, Wnt signaling, inflammatory mediator regulation of TRP channels, T cell receptor signaling, Yersinia infection, and bacterial invasion of epithelial cells also showed enrichment.

**Fig. 8.**
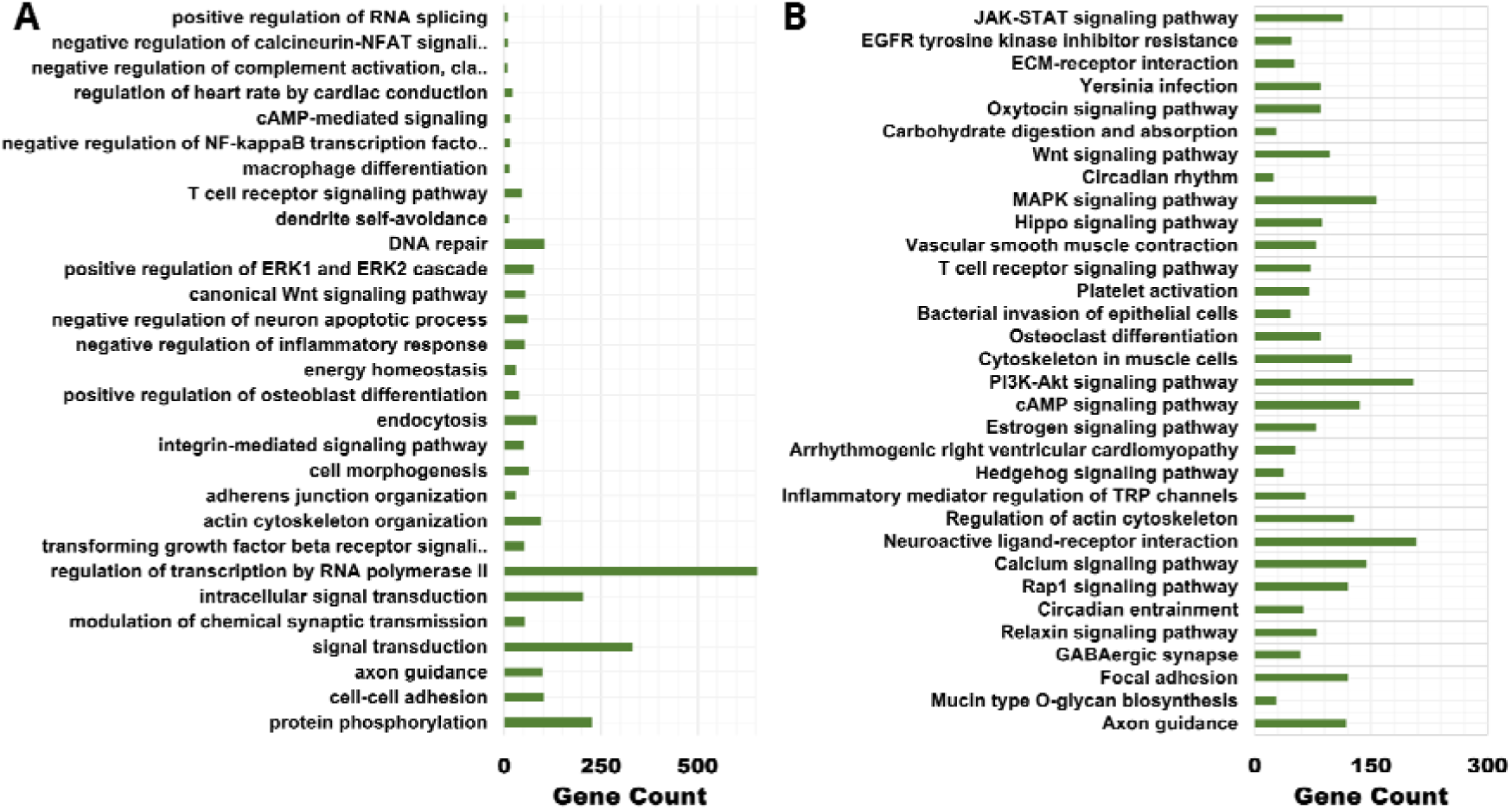
GO term and pathway enrichment for the genes had insertions in Nelore breed. The horizontal bar length indicates number of genes with variants were enriched. (A) GO term for biological process (BP) enrichment analysis result for genes with insertions. (B) KEGG pathway enrichment analysis result for the genes with insertions. These results were filtered based on P-Value < 0.05.

For genes with insertions in the Gir breed, enriched biological processes overlapped several categories found in Nelore breed while also showing distinct signatures (Fig. 9A). Enriched processes included regulation of transcription by RNA polymerase II, cell adhesion, protein phosphorylation, regulation of the ERK1/ERK2 cascade, exocytosis, and axon guidance. Immune-associated processes such as spleen development, response to bacterium, interleukin-1–mediated signaling, B cell receptor signaling, and negative regulation of NF-kappaB transcription were enriched. KEGG analysis (Fig. 9B) identified PI3K-Akt signaling, Ras and Rap1 signaling, cAMP signaling, and cell adhesion pathways. All these pathways play a role in cellular proliferation, immune modulation, and stress response. Several immune-related pathways were also prominent, including T cell receptor signaling, MAPK signaling, ErbB signaling, viral infection pathways (for example, human papillomavirus), and Fc receptor–mediated processes. These results indicate that insertions in the Gir breed associate strongly with signaling and immune pathways that may contribute to adaptability and resilience under diverse environmental conditions.

**Fig. 9.**
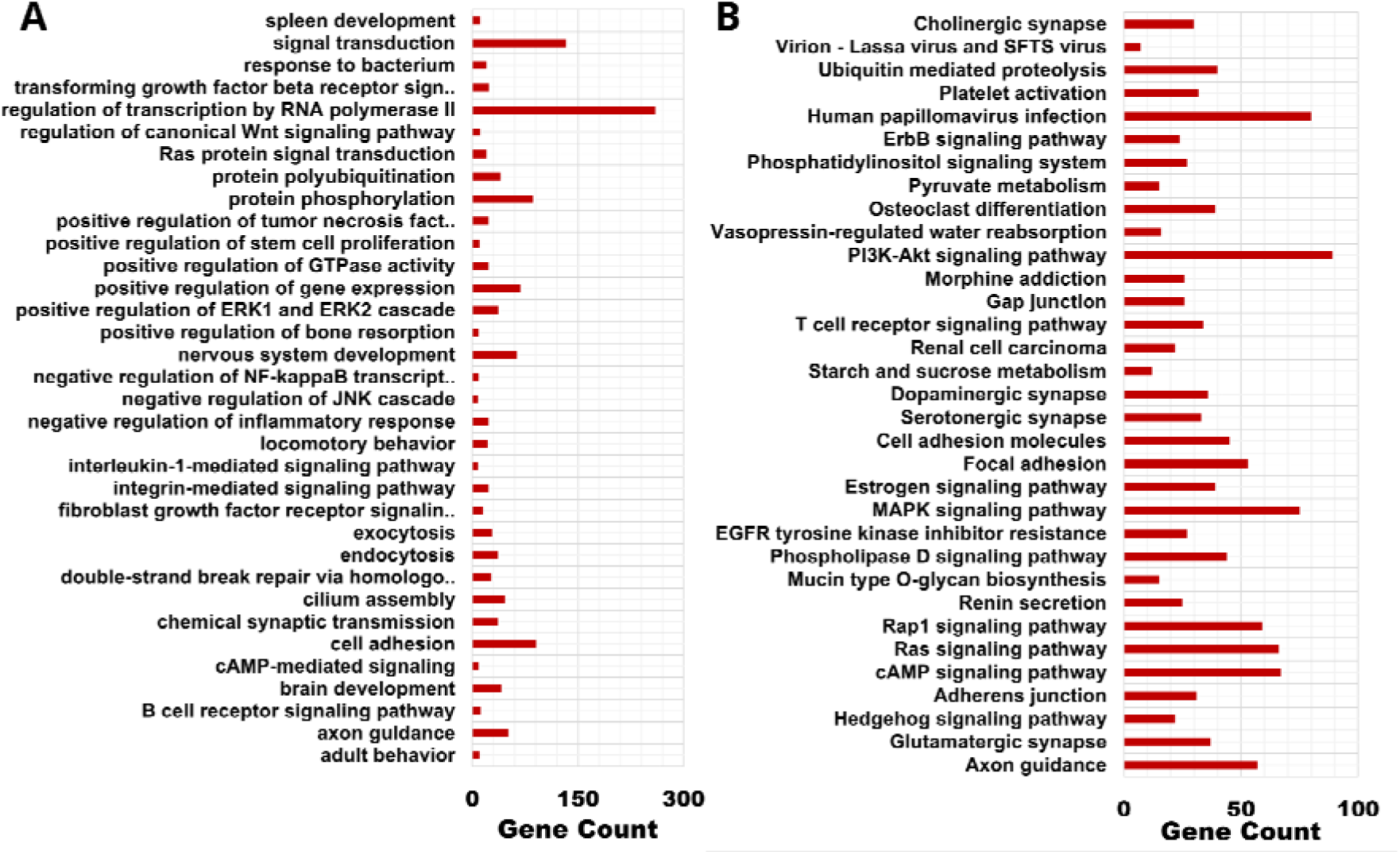
GO term and pathway enrichment for the genes had insertions in Gir breed. The horizontal bar length indicates number of genes with variants were enriched. (A) GO term for biological process (BP) enrichment analysis result for genes with insertions. (B) KEGG pathway enrichment analysis result for the genes with insertions. These results were filtered based on P-value < 0.05.

Similarly, for genes with deletions in the Nelore breed, enriched biological processes (Fig. 10A) included signal transduction, protein phosphorylation, regulation of DNA-templated transcription, ERK1/ERK2 cascade regulation, protein localization, axon guidance, and protein ubiquitination. Immune-associated processes such as regulation of inflammatory response, T cell receptor signaling, and negative regulation of NF-kappaB transcription were also enriched. KEGG pathway analysis (Fig. 10B) showed significant enrichment in PI3K-Akt signaling, Ras and Rap1 signaling, cAMP signaling, and pathways in cancer. Several immune-related pathways were enriched, including T cell receptor signaling, MAPK signaling, ErbB signaling, viral infection pathways such as human papillomavirus, and Yersinia infection.

**Fig. 10.**
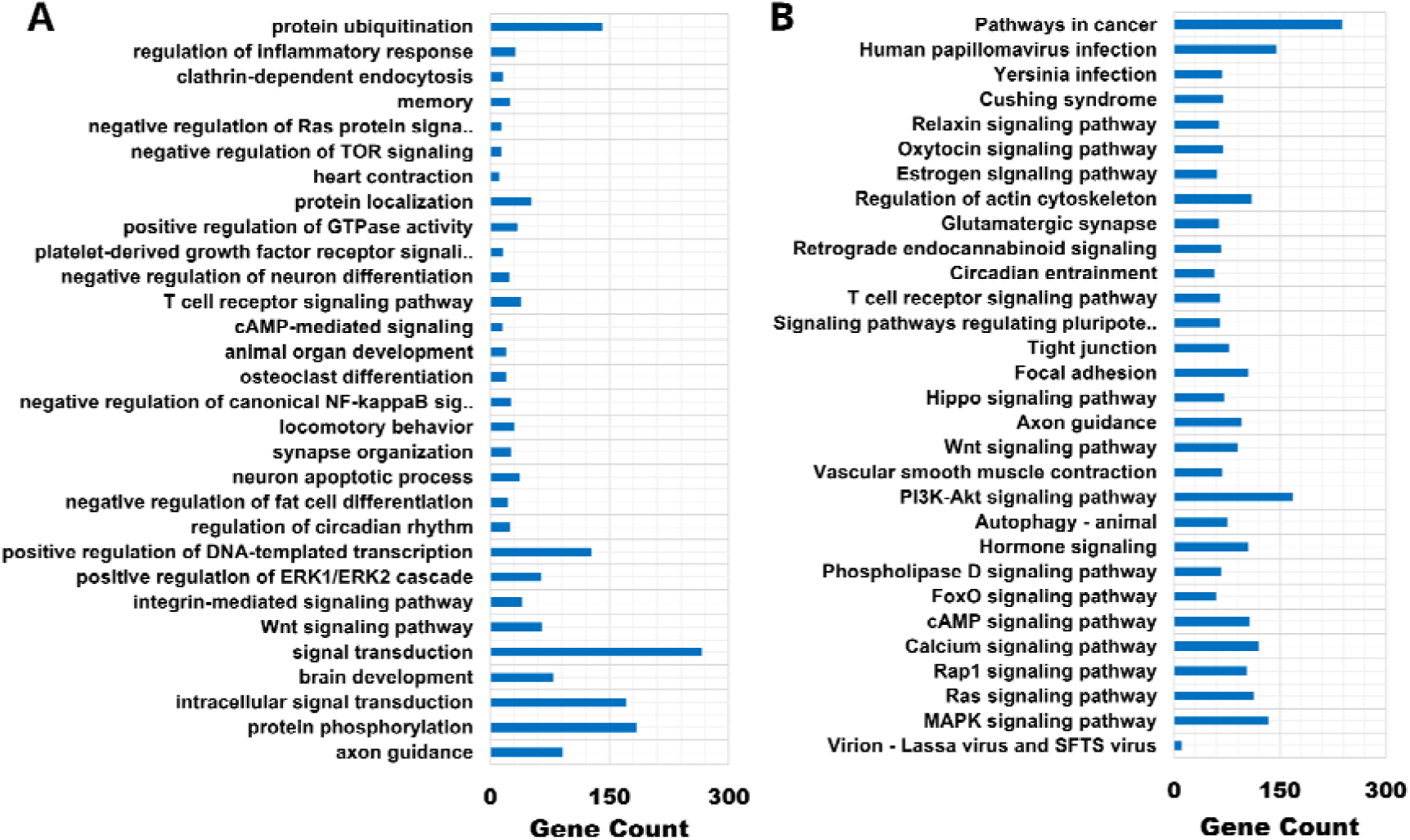
GO term and pathway enrichment for the genes had deletion in Nelore breed. The horizontal bar length indicates number of genes with variants were enriched (A) GO term for biological process (BP) enrichment analysis result for genes with deletions. (B) KEGG pathway enrichment analysis result for the genes with deletions. These results were filtered based on P-value < 0.05.

In the Gir breed, enriched biological processes (Fig. 11A) included positive regulation of transcription by RNA polymerase II, signal transduction, transmembrane transport, cell adhesion, axon guidance, protein phosphorylation, and nervous system development. Immune-associated processes such as macrophage activation involved in immune response and endocytosis were enriched. KEGG pathways (Fig. 11B) showed enrichment in PI3K-Akt signaling, calcium signaling, and immune-related pathways such as T cell receptor signaling, MAPK signaling, Fc gamma R-mediated phagocytosis, inflammatory mediator regulation of TRP channels, and Yersinia infection.

**Fig. 11.**
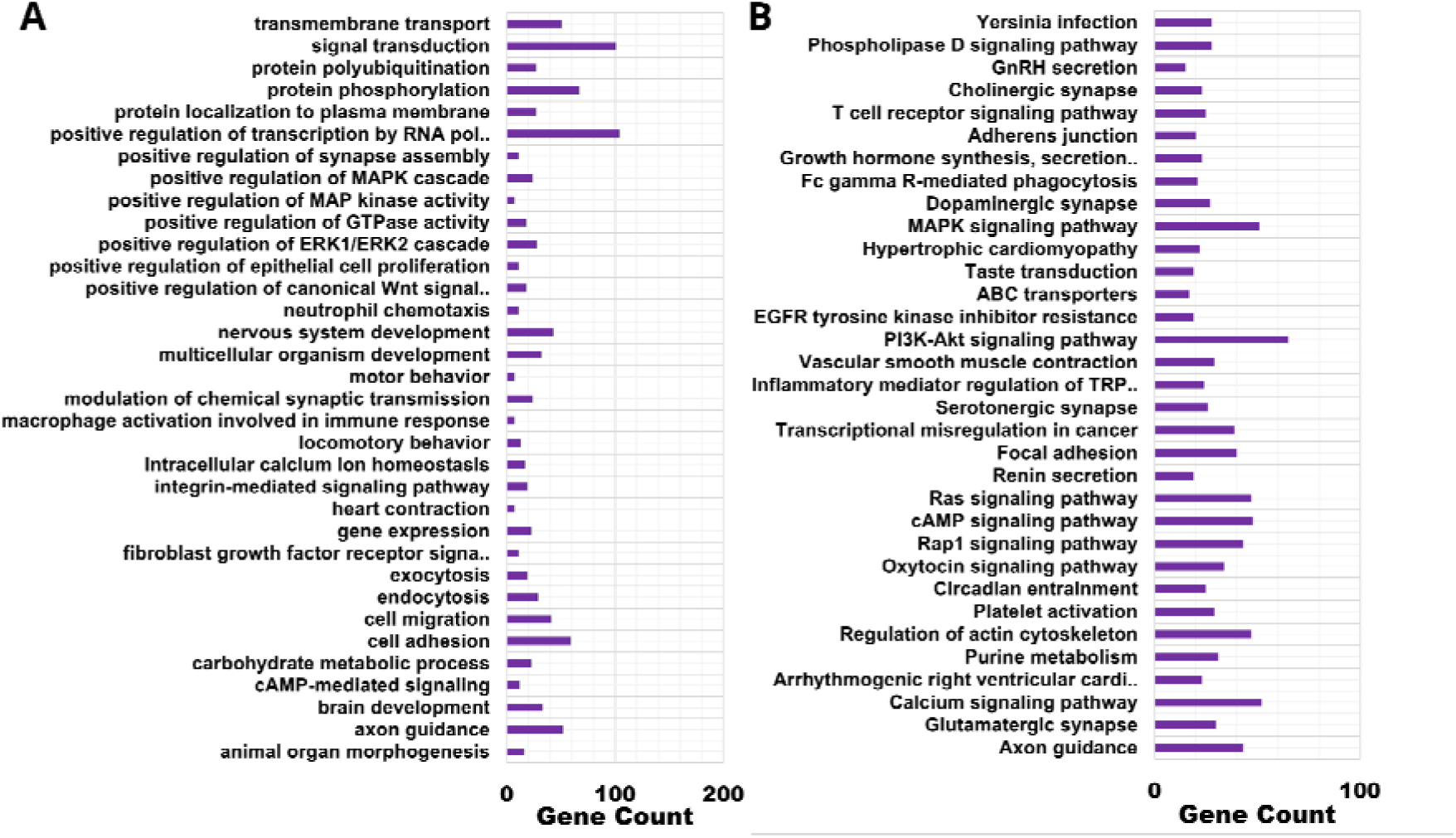
GO term and pathway enrichment for the genes had deletions in Gir breed. The horizontal bar length indicates number of genes with variants were enriched. (A) GO term for biological process (BP) enrichment analysis result for genes with deletions. (B) KEGG pathway enrichment analysis result for the genes with deletions. These results were filtered based on P-value < 0.05.

Similarly, in the Nelore breed, genes with SNVs showed enriched biological processes (Fig. 12A), including regulation of transcription by RNA polymerase II, G protein coupled receptor signaling, signal transduction, DNA damage response, ERK1/ERK2 cascade, and immune-related processes such as thymus development, regulation of autophagy, T cell activation, T cell differentiation, regulation of inflammatory response, and NF-kappaB transcription factor. KEGG pathways (Fig. 12B) included PI3K-Akt signaling, pathways in cancer, Rap1 signaling, cGMP-PKG signaling, Wnt signaling, Relaxin signaling, chemokine signaling, and thyroid hormone synthesis, and immune-related pathways such as Fc gamma R-mediated signaling, MAPK signaling, bacterial invasion of epithelial cells, and inflammatory mediator regulation of TRP channels.

**Fig. 12.**
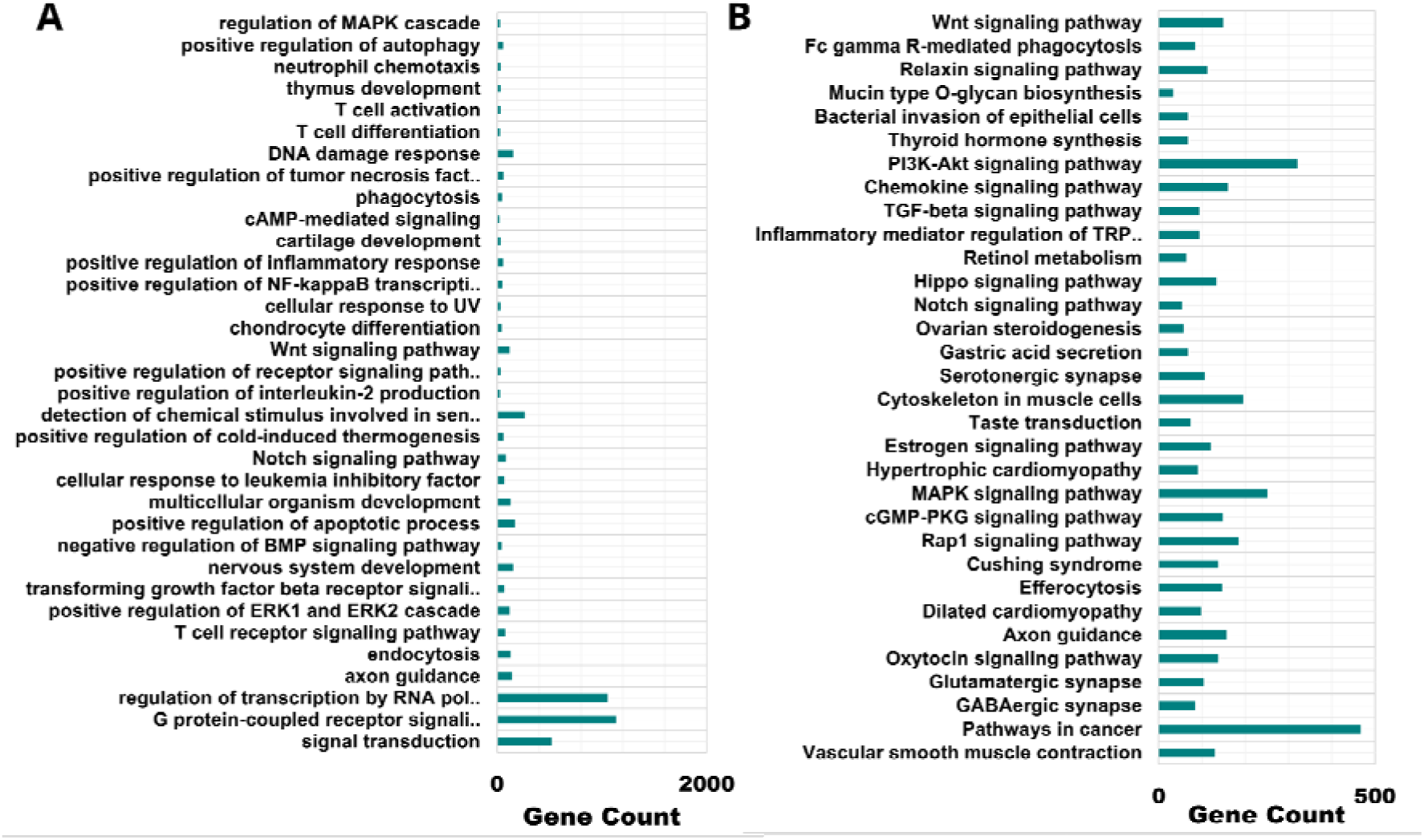
GO term and pathway enrichment for the genes had SNVs in Nelore breed. The horizontal bar length indicates the number of genes had variants were enriched. . (A) GO term for biological process (BP) enrichment analysis result for genes with SNVs. (B) KEGG pathway enrichment analysis result for the genes with SNVs. These results were filtered based on P-value < 0.05.

In the Gir breed, enriched biological processes (Fig. 13A) included G protein-coupled receptor signaling pathway, regulation of transcription by RNA polymerase II, signal transduction, protein phosphorylation, regulation of apoptotic process, regulation of ERK1/ERK2 cascade, regulation of tumor necrosis factor production, regulation of gene expression, regulation of cold-induced thermogenesis, lactation, DNA damage response, and immune-related processes such as thymus development, inflammatory response, endocytosis, phagocytosis, engulfment, and cellular response to leukemia inhibitory response. KEGG pathways (Fig. 13B) included pathways in cancer, neuroactive ligand-receptor interaction, Rap1/Ras signaling, hedgehog signaling, cAMP signaling, hormone signaling, and immune-related pathways such as NK-kappaB signaling.

**Fig. 13.**
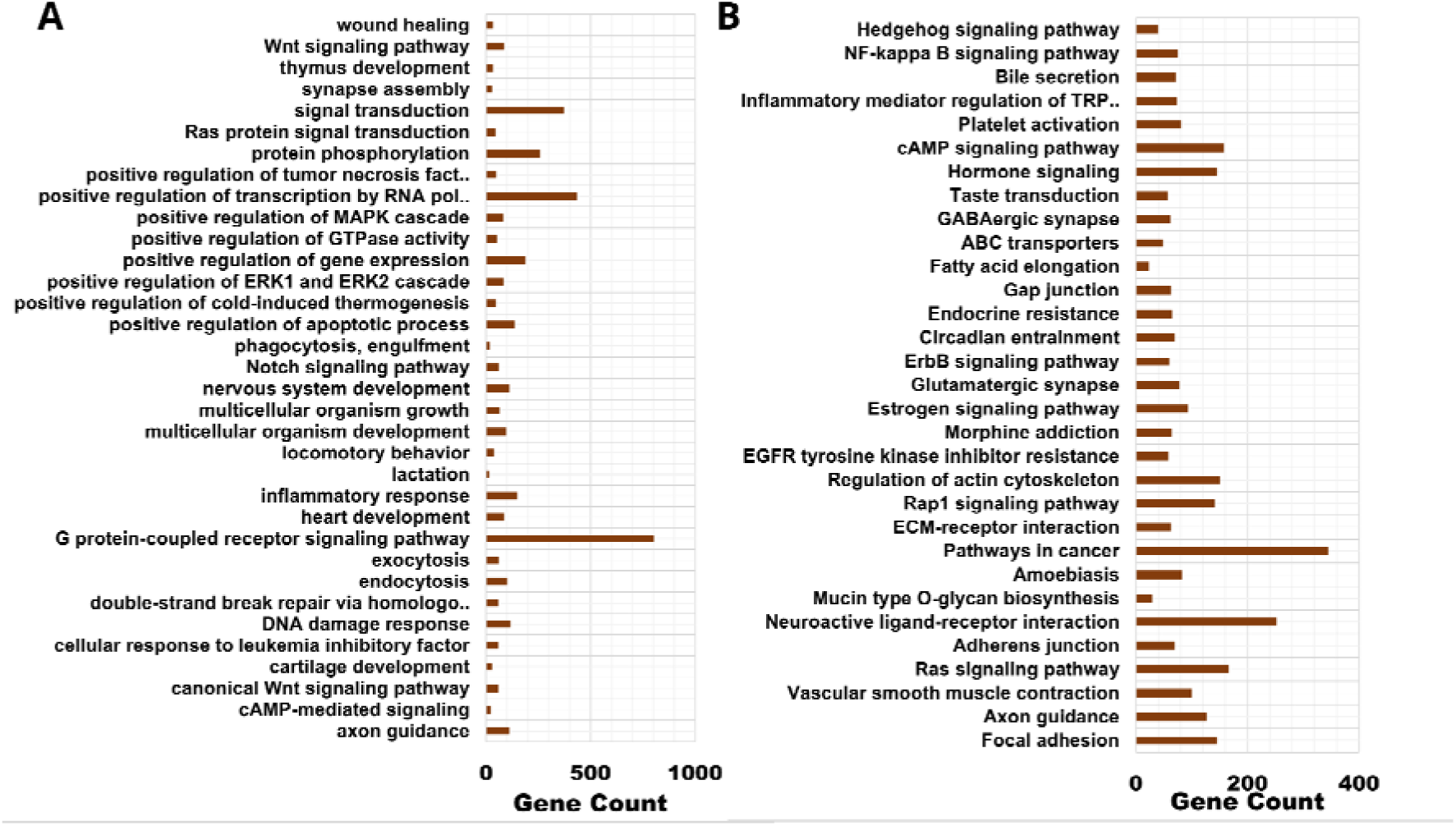
GO term and pathway enrichment for the genes had SNVs in Gir breed. The horizontal bar length indicates the number of genes had variants were enriched (A) GO term for biological process (BP) enrichment analysis result for genes with SNVs. (B) KEGG pathway enrichment analysis result for the genes with SNVs. These results were filtered based on P-value < 0.05.

## Discussion

We explored genomic variation between two *Bos indicus* breeds, Nelore and Gir, using the *Bos taurus* Hereford reference genome (ARS-UCD2.0), which comprises 1,958 contigs (29 autosomes, sex chromosomes, mitochondrial genome, and unplaced scaffolds), totals 2.77 Gb, and has an N50 of 103.3 Mb. The assembly’s high continuity and low gap content made it suitable for comparative analysis.

We performed variant calling with GATK and annotated variants with SnpEff across autosomes, sex chromosomes, the mitochondrial genome, and unplaced scaffolds. In Nelore breed, we identified 1,884,058 indels and 13,997,533 SNVs, with a transition-to-transversion (Ti/Tv) ratio of 2.26. In Gir breed, we detected 1,457,337 indels and 11,627,881 SNVs, with a Ti/Tv ratio of 2.25, supporting the reliability of SNV calls. Nelore breed carried relatively more variants than Gir breed, and SNVs were the most abundant class, followed by insertions and deletions. In both breeds, most variants localized to noncoding regions, with intergenic variants most frequent, followed by intronic, upstream, downstream, and UTR variants. Within coding and regulatory regions, insertions and deletions occurred mainly as frameshift variants, followed by splice-site, start/stop codon, transcript ablation, and in-frame (disruptive and conservative) variants, while SNVs were predominantly missense, followed by splice-site and start/stop codon variants.

We focused on genomic differences in immune-related genes identified from InnateDB and by keyword searches for additional immune-associated genes. SnpEff classified variants into four categories: high-impact (predicted to cause protein truncation), moderate-impact (may alter protein function), modifier-impact (mainly in noncoding or regulatory regions, including UTRs), and low-impact (generally functionally neutral). In the Nelore breed, we identified 2,618 high-impact, 103,309 moderate-impact, 37,416,134 modifier-impact, and 237,935 low-impact variants. In the Gir breed, we observed 2,244 high-impact, 86,542 moderate-impact, 31,005,968 modifier-impact, and 196,473 low-impact variants. As noted earlier, modifier-impact variants were the most abundant in both breeds, reflecting the prevalence of variants in noncoding and regulatory regions. For downstream analyses, we focused on immune-related genes with high-, moderate-, and modifier-impact variants.

In Nelore and Gir breeds, the immune-related genes *TLR3* and LOC508441 (CD33) carried frameshift insertions; insertions also occurred in the regulatory regions of 602 immune-related genes in Nelore breed and 219 in Gir breed. *TLR3* recognizes double-stranded RNA, it activates endosomal signaling via the adaptor *TRIF/TICAM1*, and induces type I interferons and other inflammatory cytokines. In cattle, *TLR3* expression increases after viral and parasite challenges such as *Neospora caninum* and in vaccinated animals, indicating a role in antiviral immunity and vaccine responsiveness [42, 43]. The *CD33* (LOC508441) gene encodes a Siglec-family inhibitory receptor on myeloid cells that transmits inhibitory signals through ITIM motifs to limit immune activation and maintain immune homeostasis. Although direct studies in cattle are limited, *CD33*’s conserved function across mammals suggests that genetic variation may influence inflammatory responses, pathogen tolerance, and breed-specific immune traits [44, 45].

Frameshift variants alter the reading frame of coding sequences and produce premature stop codons, truncated proteins, or loss of functional domains, thereby impairing gene function. In livestock, frameshift mutations associate with embryonic lethality, growth defects, and immune dysfunctions [46, 47]. Variants longer than 50 bp can markedly affect gene expression and protein function. For example, *DOCK4* on chromosome 4 carried a 63 bp insertion in its regulatory region in Nelore breed. Multiple insertions occurred in other immune-related genes: *TP63* had two 10 bp insertions; DDX1 had two insertions of 12 bp and 11 bp; and *PIK3R1* had three insertions (one 36 bp and two 16 bp). Multiple small insertions in promoters, enhancers, splice sites, or UTRs can change gene expression, transcript isoforms, or splicing, and thereby modulate immune responses even when the protein coding sequence remains intact [48]. *DDX1*, a DExD/H RNA helicase, participates in antiviral innate immunity by sensing dsRNA and forming complexes with *TICAM1* that stimulate type I interferon and NF-kappaB signaling. Alterations that change *DDX1* sequence or expression, including insertions in coding or regulatory regions, can alter viral RNA sensing and downstream interferon responses, potentially modifying resistance to RNA viruses [49, 50]. TP63 is an innate immune gene due to its role in maintaining epithelial integrity, a key component of barrier immunity. It regulates the proliferation and differentiation of epithelial cells in skin, rumen, respiratory, and mammary tissues. These cells act as the first line of defense against pathogens. In addition, TP63 affects immune responses by modulating cytokine and chemokine expression in epithelial cells, thereby recruiting immune cells to infection sites [51, 52].

We found 412 immune genes with insertions unique to Nelore breed and 25 unique to Gir breed. Frameshift deletions occurred in *JAM3* in Nelore breed, which also showed variants in regulatory regions of 512 immune-related genes. *JAM3*, a junctional adhesion molecule, regulates cell–cell adhesion, endothelial barrier integrity, and leukocyte migration, and thereby influences innate and adaptive immunity [53]. Long deletions (>50 bp) appeared in *PROS1, TNIP3*, and *DOCK4* in Nelore breed. *TNIP3* (TNFAIP3-interacting protein 3) interacts with A20 (*TNFAIP3*) to enhance negative regulation of NF-kappaB activation and suppress pro-inflammatory responses [54].

In Gir breed, frameshift deletions affected LOC100138951 and *PAX5* and regulatory regions of 144 immune-related genes, of which 98 are innate immune genes. *PAX5* controls B-cell lineage commitment, development, and function, and variation in *PAX5* may affect B-cell mediated immunity [40, 55]. Overall, 390 immune-related genes with deletions were unique to Nelore breed and 23 were unique to Gir breed. High-impact SNVs in Nelore breed affected genes such as *CD46* and *IL26*, with missense variants across innate immune genes including *NLRP12, MAP3K1, CD46, IL1RL2, IFNGR1, ANXA1, IFI6, MAPK8, TECPR1, ELP2, NCKAP1L*, and *ITGAM*; 382 immune genes with SNVs were unique to Nelore breed. In Gir breed, high-impact SNVs occurred in *PLG*, with missense variants in *DHX36, MARCO, IFNGR1, AKAP10*, and *DEFB7*; 58 immune genes were unique to Gir breed.

*DEFB7,* a β-defensin, functions as an antimicrobial peptide that inhibits bacteria, fungi, and viruses and recruits immune cells to infection sites [56]. *IFNGR1* encodes the ligand-binding chain of the interferon-γ receptor and mediates macrophage activation, antigen presentation, and Th1 responses; polymorphisms in cattle link to differential resistance to infections such as bovine tuberculosis [57]. *DHX36*, an ATP-dependent helicase that resolves G-quadruplex nucleic acid structures, regulates gene expression and contributes to innate immunity by sensing viral RNA and modulating interferon responses [58, 59]. The coexistence of SNVs, insertions, and deletions within the same gene, as observed for TLR3 and CD46 in the Nelore breed, suggests potential cumulative effects on transcription, splicing, mRNA stability, and protein structure that could alter biological processes [60].

Analysis of ROH islands and RAiSD-identified regions revealed strong selection signals. In Nelore breed, candidate genes included *ANKRD11* and *MAGI2* (linked to meat, carcass, milk, production, and reproduction), LOC132345096 (linked to milk), *FOXP2* (linked to meat, carcass, and reproduction), *TCF12* (linked to milk, production, and reproduction), and *ATP5PO* (linked to meat and carcass). In Gir breed, signals occurred at *MEFV* and *ORIF1*, both linked to milk traits. *ANKRD11* associates with muscle development and structure in cattle, consistent with a role in meat and carcass phenotypes [61]. *MAGI2* has been implicated in Nelore breed genomic scans for selection and has been connected to growth and weight-related traits important for beef production [62].

Genes with all three variants in both breeds were enriched in the NF-kappaB signaling pathway regulates immune responses, inflammation, and cell survival in cattle by controlling expression of pro-inflammatory cytokines such as TNF-α, IL-1β, and IL-6 and by directing activation, proliferation, and differentiation of immune cells including macrophages and dendritic cells. Variants affecting genes in this pathway, including insertions, deletions, and SNVs identified in Nelore and Gir breeds, may alter cytokine production, immune cell recruitment, and overall immune responsiveness, and thus may affect disease resistance, inflammatory regulation, and vaccine responses. These findings underscore the role of NF-kappaB in shaping breed-specific immune profiles in bovines [63–65].

We also observed enrichment of genes with variants in MAPK signaling and T cell receptor signaling pathways. Studies in bovines show that modulation of TCR signaling directly affects immune competence: TCR engagement governs T cell activation, proliferation, and cytokine secretion, which determine protective responses after infection or vaccination. Variation in TCR-mediated signaling associates with differences in vaccine responsiveness and susceptibility to respiratory pathogens in cattle [66]. Experimental infections, for example with foot-and-mouth disease virus (FMDV), demonstrated that alterations in TCR signaling and downstream cytokine networks can influence infection outcome and adaptive immune memory [67]. These observations indicate that genetic or functional variation in TCR pathways could modify both disease resistance and vaccine efficacy in cattle.

The MAPK signaling cascade transmits extracellular pathogen signals into cellular responses and regulates pro-inflammatory cytokine expression (e.g., TNF-α, IL-1β, IL-6), thus orchestrating innate defenses during bacterial and viral infections. For instance, bovine alveolar macrophages infected with Mycobacterium bovis show robust MAPK activation that controls cytokine release and shapes host–pathogen interactions [68]. Work on glucocorticoid receptor phosphorylation further indicates that MAPK pathways fine-tune inflammatory responses, balancing protective immunity with the risk of immunopathology [69]. These findings imply that MAPK signaling variants may directly affect how cattle regulate inflammation and respond to pathogens.

## Conclusion

Cattle crossbreeding in India has improved milk yield but the crossbred cattle are more susceptible to diseases. Currently, comparative genomics studies between Indicine and Taurine have not focused on disease resistance that can be used for future molecular breeding programs to improve immunity.

We performed a genome-wide comparison of two Indicine breeds (Nelore and Gir) using whole-genome alignment (BWA-MEM), variant calling (GATK HaplotypeCaller), annotation (SnpEff/SnpSift), runs-of-homozygosity analysis (PLINK), sweep detection (RAiSD), and QTL overlap using bedtools and Animal QTLdb. Nelore breed carried a larger burden of variants than Gir breed and exhibited sharper RAiSD peaks, while ROH islands highlighted breed-specific homozygous regions. Notable candidate loci under selection in Nelore breed included *ANKRD11, MAGI2, FOXP2, TCF12* and *ATP5PO*, whereas *MEFV* and *ORIF1* were found in Gir breed. At the gene level, we observed disruptive changes in immune genes such as *TLR3* and a *CD33*-like locus (LOC508441/CD46) showed frameshift events, *CD46* has a stop-gained SNVs, and multiple innate immune genes, such as *TLR3* and *CD46*, harbored combinations of SNVs, insertions, and deletions present across coding, non-coding regions, and regulatory regions. QTL mapping linked many of these variant-bearing genes to production and health traits (milk, carcass, reproduction), indicating pleiotropic associations between immune variation and other phenotypes. Functional enrichment of genes with all the three types of variants are implicated in immune signaling pathways, notably NF-kappaB, MAPK, PI3K-Akt, and T cell receptor–related processes.

These results suggest that indicine breeds show breed-specific patterns of genetic variation and selection that affect both immunity and production traits. The concentration of high-impact, moderate-impact and modifier-impact variants in immune-related genes, together with co-location at QTLs and selection peaks, prioritizes candidates for follow-up functional and association studies. Specifically, *TLR3, CD46, ANKRD11, MAGI2, LOC132345096, FOXP2, TCF12*, and *ATP5PO* in the Nelore breed, and *MEFV* and *ORIF1* in the Gir breed. However, translating these genomic signatures into breeding decisions will require independent validation: targeted genotyping in larger cohorts, expression and protein assays to test functional consequences of frameshifts and stop-gains, and controlled phenotype–genotype association analyses. Overall, the study provides a prioritized catalogue of variant-bearing and putatively selected genes that can guide future work on disease resistance and productivity in indicine cattle.

## Funding

Indian Council of Agricultural Research-National Agricultural Science Fund (F. No.NASF/SUTRA-02/2022-23/50) to RMY, DS, and SKO.

## Acknowledgment

The authors thank ICAR-National Agricultural Science Fund, SASTRA Deemed to be University, and ICAR-National Dairy Research Institute. The authors thank Mr. Pulkit Anupam Srivastava for discussions related to the Python scripts. The authors thank Prof. Dr. S. Panchapakesan, Coordinator, Central Animal Facility at SASTRA Deemed to be University for support and guidance.

## Data Availability

The datasets used in this analysis were sourced from previously submitted datasets in the NCBI database, as cited in the manuscript. The output data relevant to this study is provided within the manuscript, and all generated output files and utilized in-house Python script are available in the Zenodo repository: https://doi.org/10.5281/zenodo.17164084

## Notes

https://doi.org/10.5281/zenodo.17164084

### Competing Interest Statement

The authors have declared no competing interest.

### Summary of Updates

We have meticulously addressed all the concerns raised by the reviewers. We have revised the manuscript with a high-quality Nelore and Gir breed datasets and ensured that the scientific rigor of the manuscript and the results are not biased. Since we have taken high-quality datasets compared to the earlier data, our results are different and more reliable. While, the broad outcome remains the same, the details of the genes involved in immunity are statistically and scientifically significant.

